# Multi-omic dissection of reversible and persistent molecular alterations in diet-induced obesity

**DOI:** 10.64898/2026.09.08.749914

**Authors:** Jordi Rofes, Joan Miro-Blanch, Pau Gama-Perez, Jordi Capellades, Christian M. Heyer, Ignasi Forné, Luisa Santus, Aurélie Balvay, Claire Maudet, Sylvie Rabot, Belén Carbonetto, Pedro González-Torres, Marta Melé, Axel Imhof, Matthias Schlesner, Pablo M. Garcia-Roves, Oscar Yanes

## Abstract

Diet-induced obesity drives broad molecular remodeling across host and microbial systems, but why lifestyle intervention reverses some of these alterations while others persist remains unclear. To address this, we performed a multi-omic characterization of diet-induced obese mice after a combined nutritional and exercise intervention, integrating liver transcriptomics, epigenomics, metabolomics and metallomics with gut metagenomics and metallomics. Multi-omics factor analysis resolved two dominant axes of variation. The first captured a broadly reversible response (∼72% of altered variables), restored by dietary restriction and exercise, involving coordinated remodeling of hepatic metal homeostasis, epigenetic regulation, and immune and cell-turnover pathways. The second comprised persistent alterations resistant to intervention and driven primarily by microbial functional profiles: notably, functional diversity remained reduced despite substantial taxonomic recovery. Using germ-free mice to define the microbiota-responsive hepatic space, we found that a significant fraction of the persistent liver features fell within it, enriched in lipid metabolism (PPAR signaling, steroid and cholesterol biosynthesis, peroxisomal activity) and retinol metabolism. Sequential correlation analysis traced these changes to the loss of specific low-abundance taxa and their biosynthetic capacities, particularly vitamin (folate, biotin, cobalamin, pantothenate, thiamine) and cofactor metabolism, implicating microbiota-derived vitamin metabolism in sustained hepatic dysfunction. Cobalt was the sole essential element that remained persistently dysregulated and tracked dietary cobalt content, suggesting a diet– microbiota route to the persistent phenotype. These findings establish a dual regulatory framework in which metabolic plasticity is governed by reversible host-intrinsic and persistent microbiota-dependent processes, providing a systems-level explanation for obesogenic memory.

## Introduction

Obesity is a multifactorial disease driven by energy imbalance, unhealthy diet, and genetic predisposition^1^. Its global prevalence has doubled since 1990^2^, affecting over 16% of adults in 2022 and substantially increasing the risk of cancer, type 2 diabetes, and cardiovascular disease^3^. Despite effective lifestyle, surgical, and pharmacological interventions, including GLP-1–based therapies^4,5^, more than 80% of patients regain lost weight within five years of treatment cessation^6,7^.

Weight regain is driven by behavioral, genetic^8,9^, and physiological^10,11^ factors, but growing evidence indicates that obesity also induces persistent epigenetic remodeling in adipose tissue^12^ and the innate immune system^13^. These alterations, spanning DNA methylation changes, histone modifications, and shifts in chromatin accessibility, sustain transcriptional programs that favor lipid storage, inflammation, and reduced energy expenditure, even after high-fat diet withdrawal. This persistent “obesogenic memory” limits metabolic plasticity^14,15^ and may predispose individuals to accelerated weight regain.

The gut microbiota may provide a second layer of “obesogenic memory”^16^. Obesity induces persistent dysbiosis^17^ that is only partially reversed by lifestyle interventions^18^, with specific microbial taxa and functions remaining altered even after weight loss^19^. Such lasting microbial changes may impair metabolic plasticity and promote weight regain.

Similarly, despite excessive caloric intake, obesity is frequently associated with widespread micronutrient deficiencies^20^, including vitamins and essential minerals. Both vitamins (some derived from the gut microbiota) and minerals such as iron, copper, cobalt, zinc and selenium, influence inflammation, antioxidant defenses, and insulin regulation, with documented effects on body composition and metabolic health^21–24^. Persistent alterations in micronutrient availability may therefore reinforce obesity-associated transcriptional and metabolic programs, contributing to the maintenance of an obesogenic state.

Despite these interconnected effects, how epigenetic remodeling, gene expression, micronutrient homeostasis, and gut microbiota alterations converge to shape the persistence or reversibility of obesity remains poorly understood. The Lifestyle Matters (LiMa) project was designed to address this question by investigating how two major environmental drivers of obesity, high-fat diet and sedentarism, alter metabolic plasticity in mice, and whether intensive lifestyle intervention combining dietary restriction and exercise can reverse these effect^25^. The experimental design comprised three groups: control mice fed a standard diet, mice fed a high-fat diet (HFD), and HFD mice subsequently subjected to dietary restriction and exercise.

Here, we present a comprehensive multi-omic analysis of the LiMa model focused on the liver, a key hub of systemic energy and nutrient metabolism. By jointly interrogating epigenetic regulation, gene expression, metabolic and micronutrient status, and gut microbiota composition and function across obesity and intensive lifestyle intervention, we provide, to our knowledge, the first comprehensive characterization of these interconnected molecular layers within a single experimental framework. This unique systems-level approach enables us to distinguish persistent from reversible obesity-associated alterations and to identify candidate mechanisms underlying impaired metabolic plasticity and weight regain.

## Results

### Multi-omics factor analysis uncovers a reversible response and persistent alterations

We performed a multi-omic characterization of the LiMa model (Supplementary Figure 1A), generating bulk transcriptomics, epigenomics (DNA methylation and histone modifications), metabolomics, and metallomics data from liver tissue. To capture gut-associated changes, we additionally obtained metagenomic and metallomic profiles from colon content. In total, this approach yielded 22,377 molecular features that were considered for subsequent analyses (Supplementary File 1).

Among the analyzed layers, essential metals displayed the most extensive changes, with nearly 75% of detected metals significantly regulated in either the gut content or the liver across conditions. Transcriptomic, metabolomic, and histone modification profiles also showed widespread remodeling, with more than half of the measured variables exhibiting significant differences. In contrast, DNA methylation remained comparatively stable: only ∼0.1% of genomic CpG sites (20,426 of 20.81 million CpGs), grouped into 4,591 differentially methylated regions (DMRs), were modulated by lifestyle habits. Overall, all omic layers exhibited significant alterations, indicating broad molecular remodeling induced by the high-fat diet (HFD) and the subsequent lifestyle intervention (Supplementary Figure 1B).

To integrate the different molecular layers and identify coordinated responses, we trained a multi-omics factor analysis (MOFA) model^26^ using variables that showed significance in at least one univariate group comparison across the liver and colon datasets, which corresponds to 59.8% of the total molecular features (Supplementary File 2). To determine the optimal model complexity, we trained multiple MOFA models with increasing numbers of latent factors (Supplementary Figure 2). Based on the variance explained, we selected a final model with two latent factors. The first factor captured a reversible response, characterized by variables altered by the HFD that returned toward control levels after the lifestyle intervention (INT). In contrast, the second factor represented persistent alterations, that is, variables modified by HFD that remained dysregulated despite the intervention (Figure 1A, B). Using a variable selection approach based on MOFA model weights (see Methods and Supplementary Figure 3), we assigned 7,356 omic variables to factor 1 and 2,869 variables to factor 2. Together, these variables represent 45.7% of all measured molecular features. Among the selected variables, approximately 72% were associated with the reversible response and 28% with persistent alterations (Figure 1C), suggesting that most molecular features altered by the high-fat diet can be restored following the lifestyle intervention.

**Figure 1.**
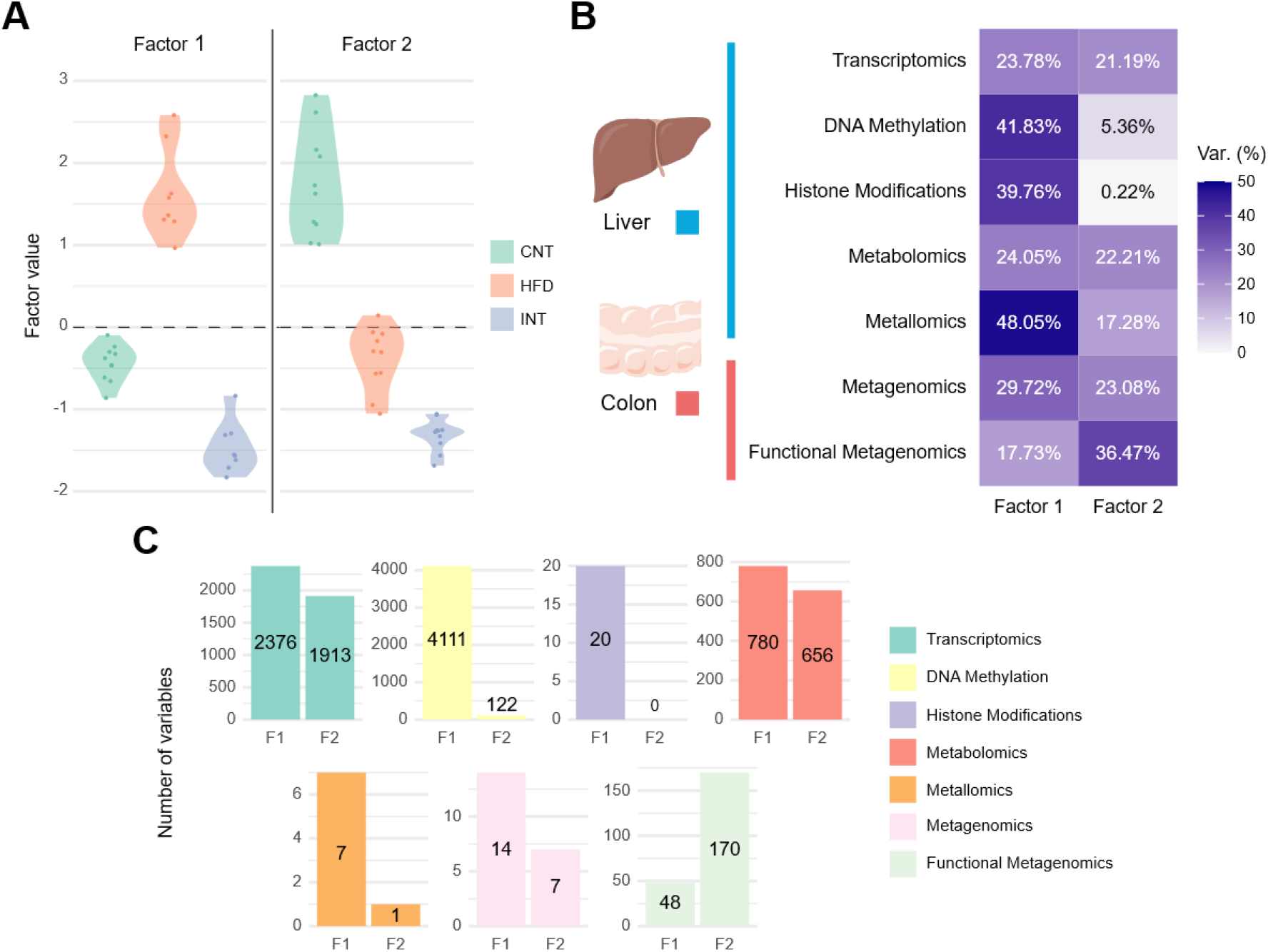
**A.** Violin plots of the two MOFA latent factors across experimental groups, illustrating reversible (Factor 1; F1) and persistent (Factor 2; F2) responses. **B.** Variance explained by the first two MOFA factors across each omic layer. **C.** Bar plots showing the distribution of variables associated with each factor after applying the selection threshold based on MOFA weights (see Methods).

Notably, a substantial proportion of the measured molecular features did not show significant changes across conditions. Out of the total of 22,377 variables profiled across all omic layers, 8,996 (40.8%) were not significantly altered by either the high-fat diet or the lifestyle intervention (Supplementary File 3). While these variables were not included in the MOFA model, their stability provides an internal reference framework of the molecular landscape that remains resilient to dietary and behavioral perturbations.

The reversible response captured by factor 1 was strongly associated with hepatic epigenetic marks, including DNA methylation and histone modifications (Figure 1B). Nearly 90% of HFD-induced differentially methylated regions (DMRs; 4,111 of 4,591) returned toward control levels following the intervention (Figure 1C). WGB sequencing revealed that HFD predominantly drives a global shift toward hypermethylation relative to controls, whereas the lifestyle intervention not only reverses this trend but shifts methylation levels below the control mean, toward a hypomethylated state, indicating an active remodeling process (Supplementary Figure 4). A similar pattern was observed for histone modifications: all 20 histone H3 modifications altered by HFD (out of 35 distinct marks detected) were restored after the intervention. Among these, modifications at H3K56, H3K79, and H3K122 exhibited the largest changes, whereas H3K9/K14 and H3K18/K23 were comparatively less affected (Supplementary Figure 5).

Consistent with the reversible molecular signature of factor 1, seven metals displayed coordinated changes in both liver and gut (Supplementary Figure 6). All seven metals showed reduced hepatic abundance under the HFD, followed by restoration toward control levels after the lifestyle intervention. Interestingly, while metal concentrations in the gut were generally strongly correlated with their dietary abundance (see concentration of minerals in mouse pellets in Supplementary Table 1), this relationship was not consistently observed in the liver. Hepatic levels of iron (Fe), copper (Cu), manganese (Mn), selenium (Se), and zinc (Zn) showed no clear association with their dietary intake, suggesting the involvement of regulatory mechanisms governing their absorption, storage, or systemic distribution.

In contrast, the persistent alterations represented by the second factor were primarily driven by microbial functional profiles, as reflected by cluster of orthologous groups (COG) annotations derived from gut metagenomic data (Figure 1B). Liver gene expression, metabolites abundance, and the taxonomic composition of the gut microbiota contributed similarly to both factors, indicating that these molecular layers participate in both reversible and persistent responses to dietary intervention.

### Coordinated epigenetic remodeling underlies the recovery of immune, cellular turnover, and metabolic pathways

Given the strong contribution of epigenetic features to factor 1, we investigated the potential regulatory impact of DMRs and histone modifications associated with the reversible response. To this end, we linked DMRs to nearby genes by considering both gene body regions (promoters, exons, introns and UTRs) and distal regions located within ±1 Mb of the transcription start site (TSS)^27^ (Supplementary File 6). Gene body DMRs were associated with 6.5% of genes whose expression was altered by HFD and subsequently restored by the lifestyle intervention (adj. p < 0.05). Notably, ∼15% of these genes encoded transcription factors (TFs) (Supplementary Table 2), significantly exceeding the proportion expected by chance among all expressed genes (7.3%; permutation test, p* < 0.05; Supplementary Figure 7A). This enrichment was, however, comparable to that observed among the narrower set of 2,376 transcripts contributing to Factor 1 (15.2%; permutation test, p < 0.05; Supplementary Figure 7B).

To evaluate whether these methylation-associated TFs could exert broad regulatory influence over the Factor 1 transcriptional program, we interrogated known TF–target relationships using the TFLink database^28^. TFLink annotations covered 90.03% of Factor 1 genes, and the combined target space of the methylation-associated TFs encompassed up to 99.39% of these genes at network distance 1, significantly exceeding random expectation (permutation test, p < 0.01; Supplementary Figure 8). Together, these findings suggest that a relatively small set of epigenetically regulated TFs may act as upstream regulatory hubs, translating DNA methylation changes into a broad and coordinated transcriptional response underlying the reversible phenotype captured by Factor 1.

Factor 2 showed a distinct regulatory architecture. Among the 16 gene body DMR-associated genes, only one TF was present (6.25%), consistent with random expectation (permutation test, p > 0.05; Supplementary Figure 7D). Broadening the analysis to the full set of Factor 2-associated genes (n = 1,913) revealed a modest but statistically significant TF enrichment (11.92%; permutation test, p < 0.05; Supplementary Figure 7F), indicating that transcription factor regulation contributes to Factor 2, albeit less prominently than in Factor 1. This is further supported by the observed-to-expected enrichment ratios: 2.04 for Factor 1 versus 1.63 for Factor 2, indicating that epigenetically driven transcriptional control is a more defining feature of the reversible response than of the persistent alterations.

Gene body DMRs were predominantly located within intronic regions and promoters (86.53%) and showed a negative correlation with the expression of their associated genes in 71.86% of cases, that is, DNA hypermethylation was associated with reduced gene expression, and vice versa. This directional bias was significantly greater than expected by chance (observed-to-expected ratio: 1.65; ***p < 1×10^-4^; permutation test; Supplementary Figure 9), confirming a non-random and directionally consistent inverse relationship between DNA methylation and gene expression at these loci.

In contrast, distal intergenic DMRs, associated with 28.8% of genes showing expression recovery, showed a more balanced distribution of positive and negative correlations, with a significantly weaker directional bias toward negative associations (observed-to-expected ratio: 1.07; *p < 0.05), consistent with the more heterogeneous and context-dependent regulatory influence of distal elements compared to proximal promoter and intronic regions^29,30^.

Histone modifications also showed coordinated regulation with DNA methylation changes (Supplementary File 7). Genes harboring gene body DMRs were significantly associated with a higher number of histone modifications than their non-DMR counterparts (***p < 0.0001; Supplementary Figure 10), as reflected by a rightward shift in the correlation count distribution. This pattern suggests that gene body DMRs do not operate in isolation but rather mark a subset of genomic loci subject to multi-layered epigenetic regulation, where DNA methylation changes co-occur with broader histone remodeling.

To investigate the functional consequences of the genes associated with the reversible response, we performed Gene Ontology (GO) enrichment analysis on the 2,376 transcripts contributing to factor 1, including those linked to gene body DMRs (Supplementary File 8). Enriched terms were primarily related to immune system processes, ribosome biogenesis, apoptosis, and cell–cell interactions, highlighting coordinated regulation of inflammatory and cellular turnover pathways. Notably, chromatin remodeling was also significantly enriched, consistent with the prominent contribution of epigenetic regulation within factor 1 (Figure 2A).

**Figure 2.**
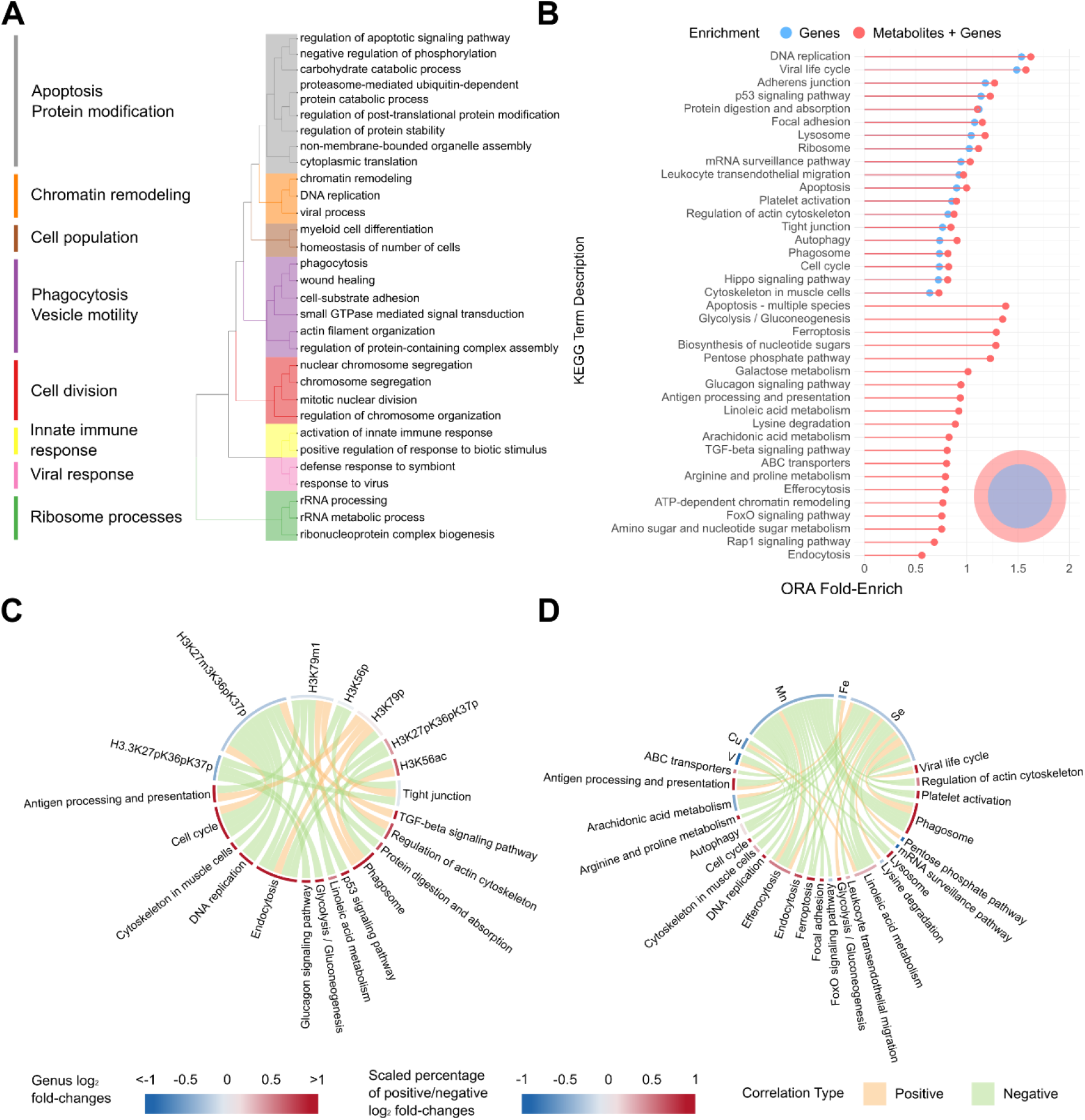
**A.** Gene Ontology (GO) enrichment cluster plot showing the top 30 overrepresented GO terms associated with Factor 1–linked genes. **B.** Lollipop plot showing log2 fold-enrichment of KEGG pathways associated with Factor 1–linked metabolites. Results are shown for gene-only enrichment (blue) and combined gene+metabolite enrichment (red). The Venn diagram indicates the overlap between KEGG terms identified in both analyses. **C.** Chord diagram showing significant correlations between histone modifications and liver KEGG pathway enrichments derived from DMR-associated genes (Pearson r ≥ 0.8, adjusted p ≤ 0.05). Edge color denotes correlation direction (positive or negative). Node colors represent HFD vs. control log2 fold-change for histone marks and the scaled proportion of positively or negatively changing genes within each KEGG pathway. **D.** Chord diagram showing significant correlations between Factor 1–associated metals and liver KEGG pathway variables. Node colors represent log2 fold-changes for metals and the scaled proportion of positive or negative gene-level log2 fold-changes within each pathway. Edge colors indicate correlation direction (orange, positive; green, negative).

To further contextualize these transcriptional changes, we integrated metabolomic features associated with factor 1 through KEGG pathway enrichment analysis (Supplementary File 9). This multi-layer approach reinforced the GO-derived signatures, particularly those related to cell–cell communication, cell death and clearance, and inflammatory processes, while expanding the functional landscape to include carbohydrate, nucleotide, and fatty acid metabolism. In addition, key regulatory pathways such as FoxO, AMPK, Rap1, and glucagon signaling, as well as core metabolic processes including glycolysis and gluconeogenesis, were significantly enriched (Figure 2B). These pathways were strongly associated (Pearson r ≥ 0.8; adj. p ≤ 0.05) with coordinated changes in histone H3 modifications, characterized by decreased methylation at H3K27 and H3K79, and increased phosphorylation at these residues under the HFD, followed by restoration after the lifestyle intervention (Figure 2C). Together, these results link reversible epigenetic remodeling to coordinated regulation of immune and metabolic pathways, supporting a central role for chromatin dynamics in the recovery of metabolic plasticity.

Additionally, correlation analysis between the seven reversible metals and KEGG pathways revealed strong links between micronutrient availability and the functional programs underlying the reversible response (Supplementary File 10). Selenium (Se) and manganese (Mn) exhibited the highest number of negative correlations with these pathways, followed by copper (Cu) and iron (Fe), whereas vanadium (V) was predominantly associated with positive correlations (Figure 2D). Among the most strongly associated pathways were efferocytosis, autophagy, and apoptosis, processes closely linked to cellular stress responses and forms of metal-dependent cell death. These associations suggest that the reduced hepatic levels of Se and Mn under the high-fat diet may contribute to the activation of stress and clearance pathways, which are subsequently normalized upon restoration of metal homeostasis during the lifestyle intervention.

### Functional microbiota remodeling underlies persistent hepatic metabolic alterations

Given the strong contribution of metagenomic variables to factor 2, we investigated the microbiota-associated persistent alterations. The shotgun metagenomic profiles from both cecum and colon samples enabled assessment of taxonomic composition and functional potential through Cluster of Orthologous Groups (COGs). At the genus level, alpha diversity (Shannon index) confirmed that the high-fat diet reduced microbial diversity in both gut regions, with a more pronounced decrease in the colon. This reduction was largely reversed by the lifestyle intervention (Figure 3A), consistent with previous reports^31,32^.

**Figure 3.**
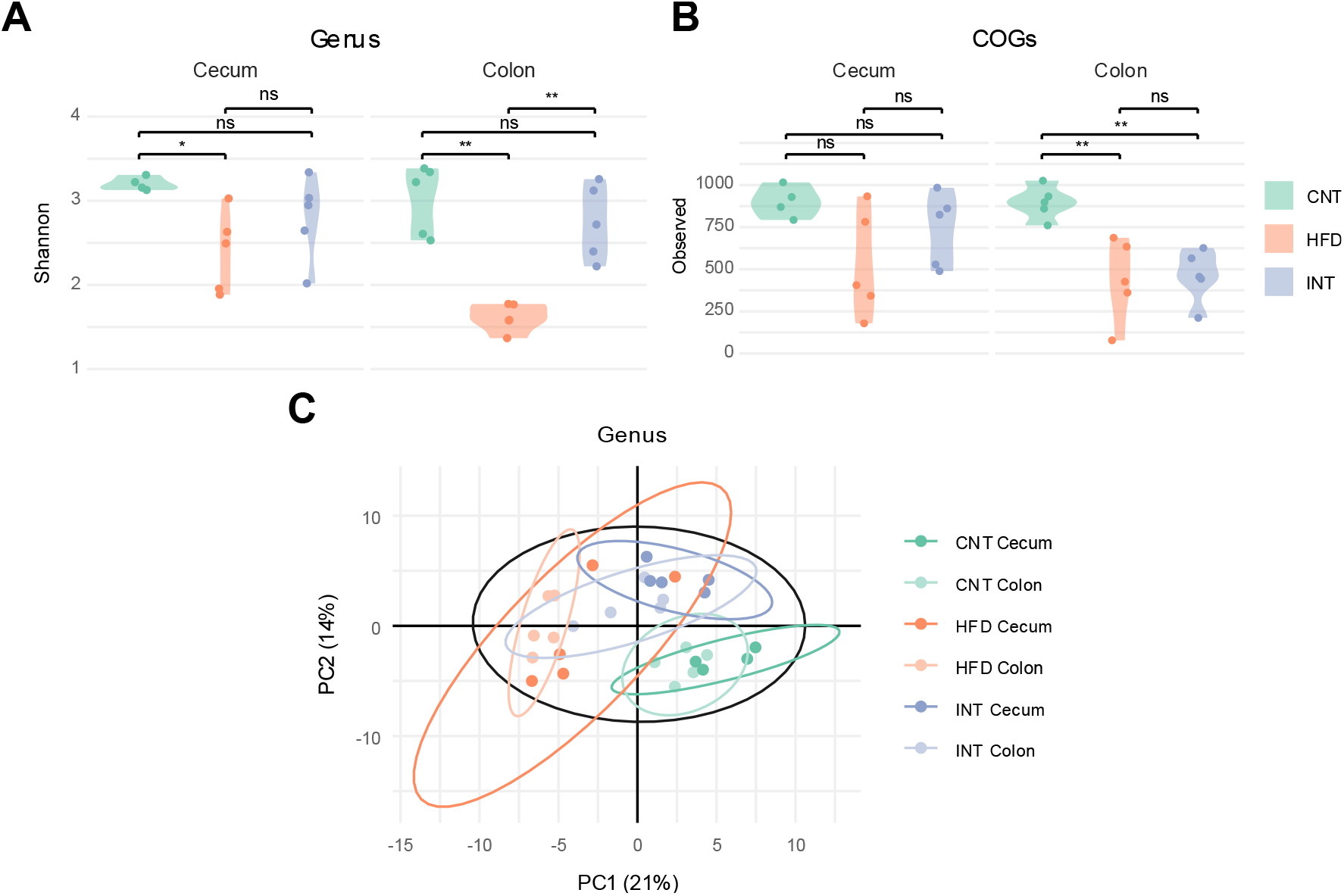
Taxonomic recovery but persistent functional impairment of the gut microbiota after lifestyle intervention. **A.** Alpha diversity (Shannon entropy) of gut microbial genera in cecum and colon across experimental groups (CNT, HFD, INT), showing HFD-induced diversity loss that is largely reversed by the intervention. **B.** Absolute abundance of microbial functional profiles (Clusters of Orthologous Groups, COGs) in cecum and colon across experimental groups, showing a functional reduction in the colon that persists after intervention. Statistical significance in A and B: *p ≤ 0.05; **p ≤ 0.01; ns, not significant (p ≥ 0.05). **C.** Beta diversity based on Aitchison distances, visualized by principal coordinates analysis (PCoA), for cecum and colon samples across experimental groups. ADONIS test results: gut region (p ≤ 0.05), diet (p ≤ 0.05), region x diet interaction (ns, p ≥ 0.05).

In contrast, microbial functional diversity exhibited a distinct pattern. The number of observed COG functionalities was significantly reduced by HFD in the colon (**p ≤ 0.01) and remained decreased after the intervention (**p ≤ 0.01) (Figure 3B). Beta diversity analysis confirmed significant effects of both dietary condition and gut region on microbial community structure (ADONIS, *p ≤ 0.05), with no significant interaction between them, indicating that dietary effects on microbiota composition are consistent across gut regions but more pronounced in the colon (Figure 3C). The cecum showed greater resilience to both HFD and intervention, with non-significant changes in functional diversity. Therefore, subsequent analyses focused on the colon, where taxonomic and functional alterations were most pronounced.

The MOFA model and diversity analyses revealed a dissociation between microbial abundance and functional importance, suggesting that, although microbial diversity may be partially restored, specific taxa carrying key metabolic functions may fail to re-establish, leading to persistent functional deficits. The 14 reversible genera (Factor 1) were numerically dominant (32.4% of the control colonic microbiome, including *Lactobacillus, Bacteroides*, *Odoribacter*, and *Roseburia*) (Supplementary Table 3) but functionally narrow, largely RNA-processing, contributing only 48 COG terms across 17 pathways (Supplementary Table 4). In contrast, the 7 persistently altered genera (Factor 2) represented just ∼4% of the colonic microbiome (Supplementary Table 5) yet encoded a disproportionately rich functional repertoire of 170 COG terms across 34 pathways, mainly involved in vitamin and cofactor biosynthesis (Supplementary Table 6).

To investigate whether persistent alterations captured by factor 2 are linked to microbiota-dependent regulation of liver functions, we employed a two-step strategy. The underlying rationale is that hepatic functions unresponsive to the presence or absence of microbiota are, by definition, unlikely to be regulated by changes in microbial community composition, such as the dysbiosis induced by HFD. Conversely, hepatic pathways that are sensitive to microbial colonization represent the biologically plausible space within which gut microbiota remodeling could exert downstream effects on liver physiology. We therefore first defined this microbiota-responsive hepatic space by comparing liver transcriptomic and metabolomic profiles between germ-free (GF) mice and conventionally raised controls (Supplementary Figure 11A). This comparison established which hepatic functions are regulated by microbiota activity. In a second step, we tested whether the liver-associated features contributing to factor 2 in the LiMa model were enriched within this microbiota-responsive space, effectively using it as a filter to assess the plausibility of a gut-liver axis driving the persistent alterations (Supplementary Figure 11B). Applying this filter, the liver-associated features of Factor 2 showed substantial and statistically significant overlap with the microbiota-responsive hepatic space: 32.8% of Factor 2 genes (627 of 1,913) and 19.4% of metabolite features (127) fell within microbiota-regulated hepatic functions, both well beyond chance expectation (permutation test: p < 1×10⁻¹⁰ for genes, p < 3×10⁻⁷ for metabolites; Supplementary Figures 11C, D and Supplementary File 11). This convergence indicates that a substantial fraction of the persistent hepatic alterations induced by HFD corresponds to functions intrinsically dependent on microbial colonization, and are therefore plausibly driven or maintained by the dysbiosis captured in Factor 2.

Gene Ontology (GO) enrichment analysis of the overlapping gene and metabolite sets revealed processes related to oxidative stress response, lipid storage and metabolism, and small-molecule metabolic pathways (Figure 4A and Supplementary File 12). Integration of metabolomic features through KEGG pathway enrichment further refined these signatures, pointing to a strong enrichment in lipid metabolism, including PPAR signaling, steroid hormone biosynthesis, unsaturated fatty acid metabolism, fat digestion and absorption, peroxisomal activity, and cholesterol metabolism (Figure 4B and Supplementary File 13). Alongside these, regulation of the circadian rhythm and pathways linked to microbiota-derived metabolites were prominently represented, including propanoate metabolism and bile secretion, as well as vitamin-related processes such as retinol metabolism, one-carbon and amino acid metabolism.

**Figure 4.**
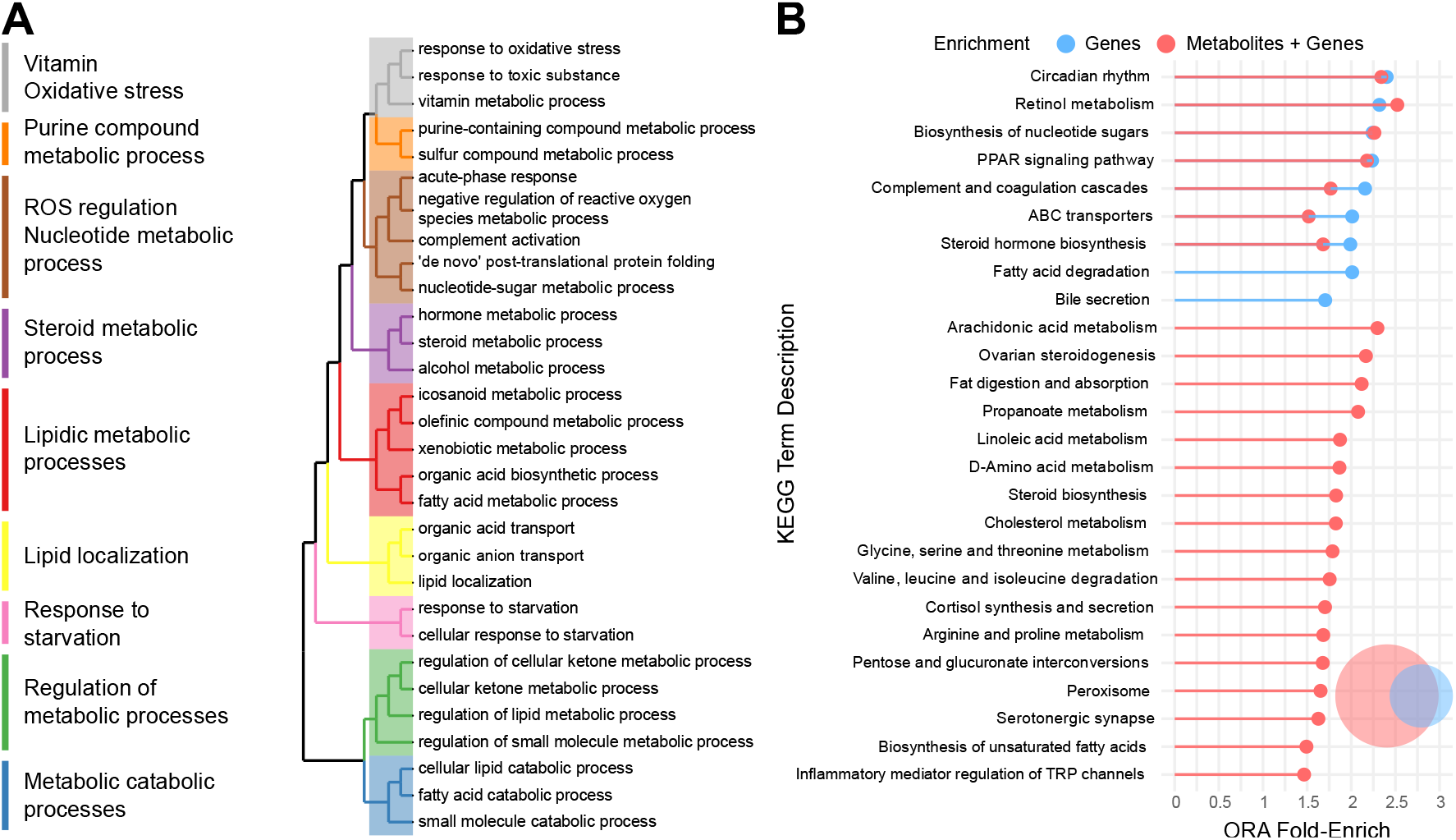
**A.** Gene Ontology (GO) over-representation analysis (ORA) of microbiota-regulated genes overlapped with factor 2. The plot shows the top 30 enriched terms grouped according to functional similarity. **B.** KEGG pathway over-representation analysis integrating factor 2-associated microbiota-regulated genes and metabolites. The lollipop plot displays the log2 fold-enrichment of pathways identified using gene data alone (blue) or the combined gene–metabolite dataset (red). The Venn diagram indicates the overlap between pathways enriched in gene-only and integrated analyses.

To dissect the microbial mechanisms underlying this persistent liver phenotype, we performed a sequential correlation analysis linking bacterial genera, their associated functional profiles, and the hepatic KEGG pathways identified above (Figure 5A and Supplementary File 14). HFD was associated with depletion of several genera, including *Parasutterella*, *Eisenbergiella*, *Anaeroplasma*, *Marvinbryantia*, *Acetatifactor*, and *Ruminococcus*, and with increased relative abundance of Lactococcus. Although these genera collectively accounted for only ∼4% of total microbial abundance in control animals (Supplementary Table 5), all showed at least one strong correlation with their corresponding COG functional profiles (Pearson r ≥ 0.75, adj. p *≤* 0.05).

**Figure 5.**
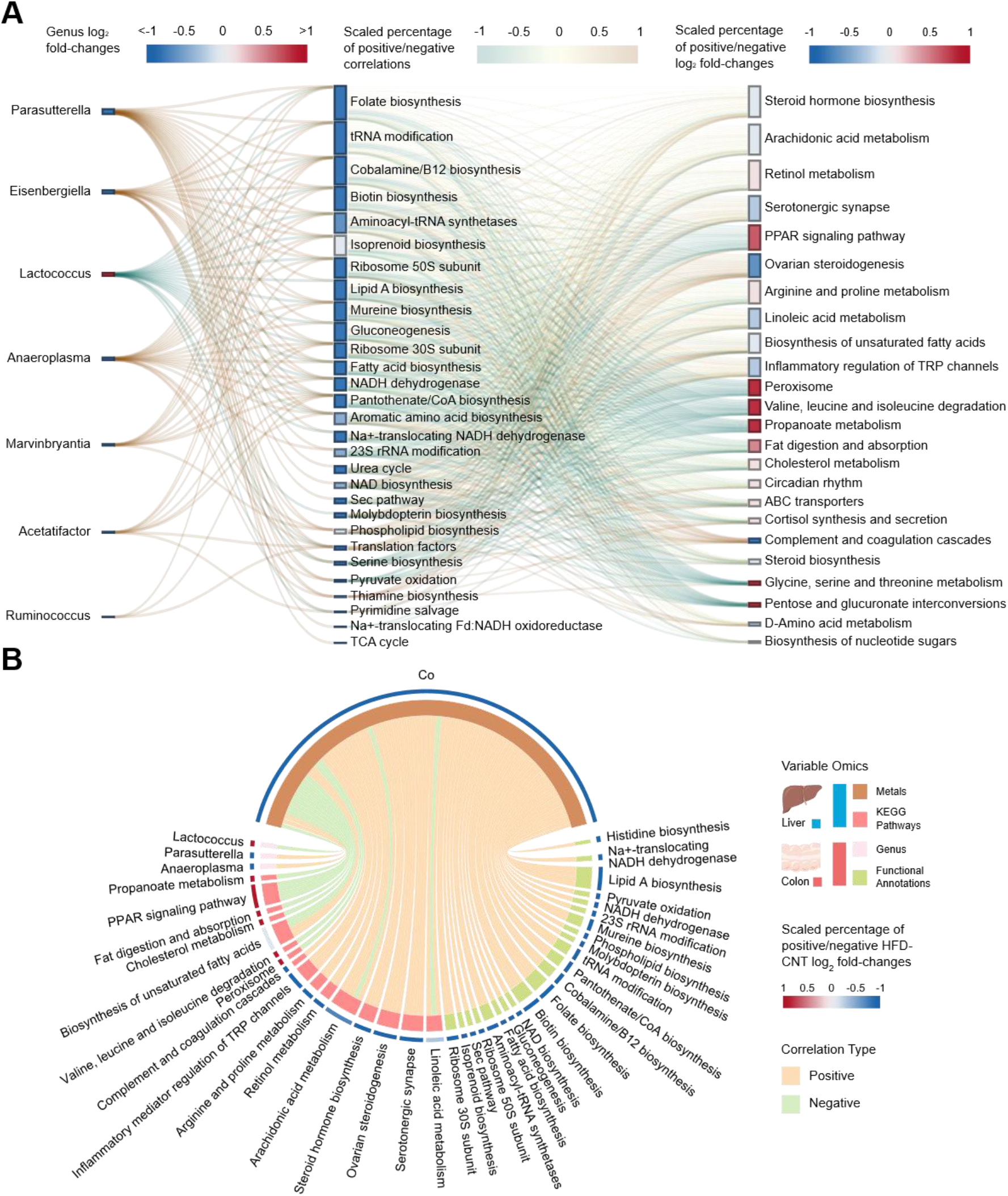
Correlation structure linking microbial taxa, microbial functions, and hepatic pathways associated with Factor 2. **A.** Sankey diagram of the strongest significant correlations (Pearson r ≥ 0.75, adjusted p ≤ 0.05) among microbial genera, microbial COG (cluster of orthologous groups) functional categories, and liver KEGG pathways associated with Factor 2. COG categories are grouped into higher-order functional annotations for visualisation. Edge colour indicates the predominant direction of correlation, from negative (blue) to positive (red). Node colour represents, for microbial genera, the HFD-versus-control log₂ fold-change, and for microbial functions and KEGG pathways, the scaled proportion of positively versus negatively altered features within each. **B.** Chord diagram of significant correlations (Pearson r ≥ 0.75, adjusted p ≤ 0.05) between hepatic cobalt levels and Factor 2-associated KEGG pathways, microbial functions, and bacterial genera. Edge colour indicates the direction of correlation (positive or negative). Node colour is encoded as in (A): HFD-versus-control log₂ fold-change for cobalt and bacterial genera, and the scaled proportion of positively versus negatively altered features for KEGG pathways and microbial functional categories.

The functional capacities most strongly associated with these genera were enriched in vitamin biosynthesis, including folate, biotin, cobalamin, pantothenate, and thiamine, as well as lipid-related processes such as fatty acid, phospholipid, and lipid A biosynthesis, and cofactor metabolism including molybdopterin and NAD biosynthesis. The HFD-induced loss of these microbial functional signatures was, in turn, strongly correlated with persistent alterations in host hepatic metabolism, particularly in lipid pathways (steroid biosynthesis, PPAR signaling, arachidonic acid, peroxisome and propanoate metabolism) and retinol metabolism (Figure 5A).

An interesting observation within the gut–liver axis involves cobalt (Co), the only essential metal associated with Factor 2. Co levels remained persistently low in both the gut and liver following the lifestyle intervention and closely tracked dietary Co intake, which differed between the control diet (66 ng/g) and the HFD and intervention diets (31 ng/g) (Supplementary Table 1). The dietary dependence of Co levels is further supported by the germ-free versus conventional comparison: despite identical dietary Co intake, gut and liver Co levels did not differ according to microbiota status, indicating that the microbiota does not determine total Co levels (Supplementary Figure 12). This pattern contrasts with the reversible metals associated with Factor 1 (Cu, Fe, Mn, Se, and V), whose hepatic levels did not correlate with dietary content in either the LiMa model or the germ-free versus conventional comparison (Supplementary Figures 7 and 12), suggesting that their availability is instead governed by host absorption and excretion, potentially modulated by microbiota composition.

Co levels were positively correlated (Pearson r ≥ 0.75, adjusted p ≤ 0.05) with microbial functional capacities for vitamin biosynthesis (folate, biotin, cobalamin, and pantothenate/CoA), lipid metabolism, and cofactor production (molybdopterin and NAD biosynthesis). Concomitantly, Co levels were linked to persistent changes in host liver metabolism, including downregulation of retinol metabolism, arachidonic acid metabolism, and steroid biosynthesis, alongside upregulation of pathways such as propanoate metabolism, cholesterol metabolism, PPAR signaling, branched-chain amino acid degradation, and peroxisomal activity (Figure 5B and Supplementary File 15).

Beyond the microbiota-driven component of Factor 2, a small but coherent epigenetic signature was identified: 16 gene body DMRs representing loci where HFD-induced methylation changes persisted despite the lifestyle intervention (Supplementary Table 7). Despite their small number, these genes converge functionally on lipid and carbohydrate catabolism. Peroxisomal fatty acid β-oxidation is represented by *Acaa1b* and *Ech1*, consistent with the persistent upregulation of peroxisomal activity identified in the KEGG enrichment analysis; lysosomal lipid catabolism by *Gm2a* and *Plbd1*; hepatic lipid transport and membrane dynamics by *Mfsd2a* and *Gpc1*; and carbohydrate and intermediary metabolism by *Mlxipl* (ChREBP), a glucose-responsive transcription factor coordinating glycolytic and lipogenic gene expression, and *Pnldc1*. Interestingly, eight of these 16 genes (*Gpc1*, *Mfsd2a*, *Tmem184a*, *Plbd1*, *Tenm3*, *Acaa1b*, *Gm2a*, and *Pnldc1*) fell within the microbiota-responsive hepatic space, as they were also differentially expressed in conventionally raised versus germ-free mice, with most showing increased expression under HFD, INT, and GF conditions. A particularly compelling example is Gm2a, which was recently reported to be hypermethylated in children with obesity^33^. Its gene body was also differentially methylated between conventionally raised and germ-free mice, and hypermethylation across HFD, INT, and GF conditions strongly correlated with reduced Gm2a expression relative to conventional controls (Supplementary Figure 13). Together, these findings suggest that the gut microbiota normally maintains the methylation state of these loci, and that HFD-induced dysbiosis disrupts this control, leaving a persistent hypermethylation mark that lifestyle intervention cannot reverse.

### Low-responder mice reveal early microbial and hepatic alterations preceding the full obese phenotype

A subset of HFD-exposed mice (n = 6) was excluded from the HFD group as they displayed partial resistance to the full metabolic obese phenotype: despite 16 weeks of identical hypercaloric exposure, they exhibited significantly lower body weights, liver weights indistinguishable from controls, and attenuated glucose intolerance and insulin resistance (Supplementary Figure 14). Rather than discarding this heterogeneity, we reasoned that these low-responders represent a naturally occurring intermediate state along the obesogenic trajectory, in which molecular alterations already present reflect early upstream events, while those absent require full phenotypic deterioration to emerge.

To test this, we trained a second MOFA model replacing the intervention group with the low-responder mice (HFDex). This model again resolved two dominant factors mirroring the original structure: Factor 1 was associated with the low-responder phenotype and driven by epigenetic and metallomic variables, while Factor 2 captured alterations shared with the full HFD phenotype and was primarily driven by microbial functional profiles (Supplementary Figure 15). Comparing the variables assigned to each factor across both models (Supplementary Figure 16) revealed a key asymmetry: Factor 1 variables overlapped by ∼80% between models (5,872 shared variables; permutation test, p < 0.0001), whereas Factor 2 variables overlapped by only ∼50% (1,451 shared variables; p < 0.0001) (Supplementary Figure 17). Although both overlaps far exceeded chance, their difference is biologically informative. The high Factor 1 overlap indicates that most of the reversible epigenetic and metallomic changes are also present in low-responders, albeit in attenuated form, consistent with these animals representing a milder, intermediate stage of the phenotype. In contrast, the partial Factor 2 overlap indicates that half of the persistent HFD-induced alterations are already established in low-responders, before the full obese phenotype develops (Supplementary File 16).

These early alterations encompassed hepatic dysregulation of circadian rhythm, PPAR signaling, and branched-chain amino acid, propanoate, retinol, taurine, hypotaurine, and lipid metabolism (Figure 6A and Supplementary Files 17, 18), several of which (PPAR signaling, propanoate, and retinol metabolism) were identified as microbiota-dependent in the original MOFA model. Consistent with this, cobalt depletion and reduced microbial biosynthetic capacity for vitamins (thiamine, pantothenate/CoA, cobalamin, folate, biotin) and cofactors (molybdopterin, NAD) were already evident in low-responders, along with dysregulation of related hepatic metabolites (Figure 6B). Together, these results indicate that microbial functional erosion and its hepatic consequences are among the earliest molecular events in diet-induced metabolic dysfunction, arising before, and potentially contributing to, full phenotypic conversion.

**Figure 6.**
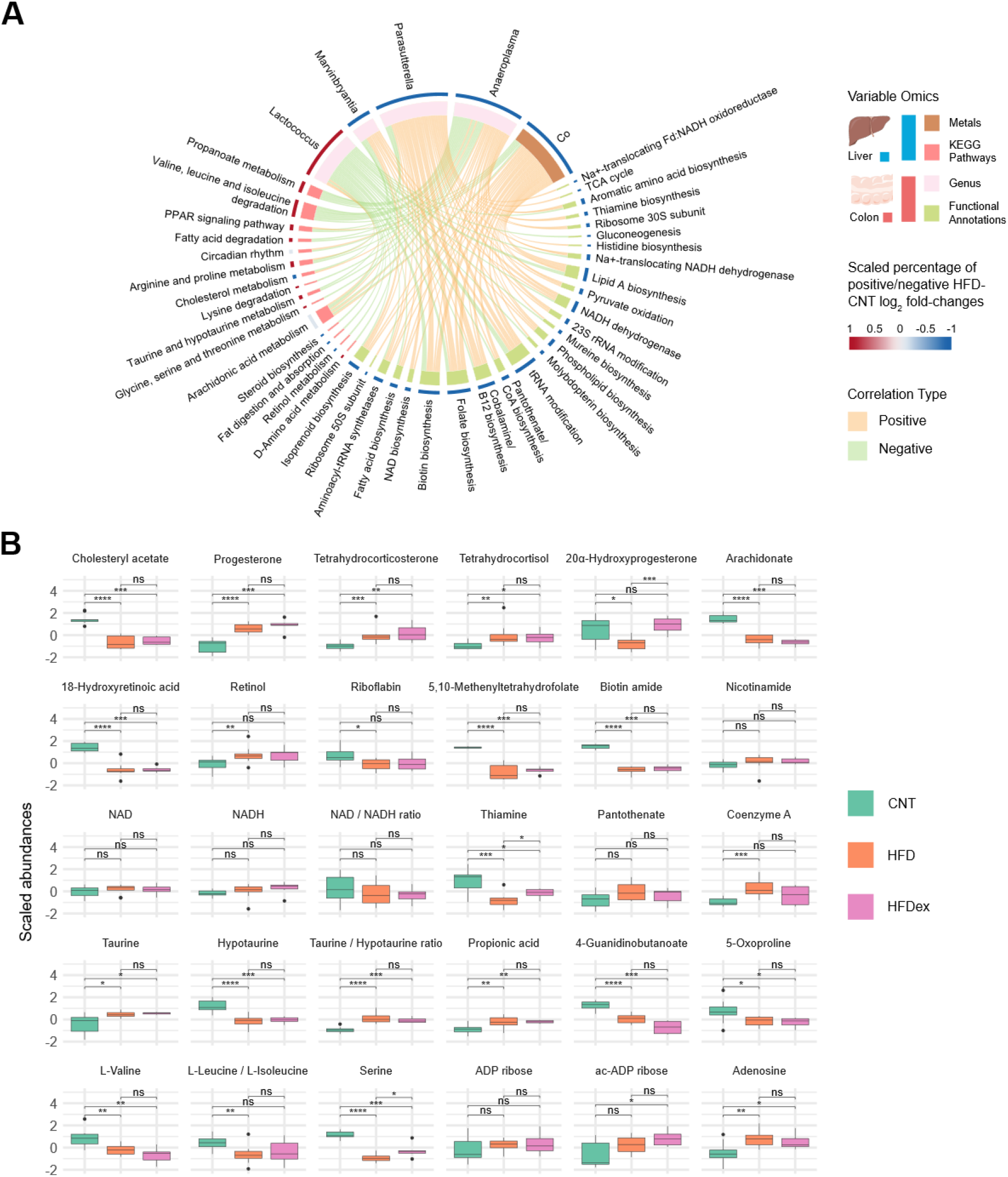
Early and persistent molecular alterations identified in low-responder (HFDex) mice. **A.** Chord diagram showing strong, significant correlations (Pearson r ≥ 0.75, adjusted p ≤ 0.05) linking colon bacterial genera, hepatic Co levels, hepatic KEGG pathways, and colon microbiota functional categories that were associated with Factor 2 in both the primary (lifestyle intervention, INT) and low-responder (HFD-excluded, HFDex) MOFA models. Nodes are grouped by omic layer and tissue: liver metals and KEGG pathways (blue arc) and colon genera and functional annotations (red arc), as indicated in the legend. Ribbon (edge) color denotes correlation direction (orange, positive; green, negative). Node color denotes the scaled percentage of positive versus negative HFD-versus-control log2 fold-changes: for bacterial genera and metals, this reflects the direction and magnitude of their own change; for KEGG pathways and functional categories, it reflects the scaled proportion of positively or negatively altered features contributing to each term (red, predominantly increased; blue, predominantly decreased). **B.** Hepatic abundances of representative metabolites across experimental groups (CNT, HFD, HFDex) for pathways already dysregulated in low-responders, spanning steroid and cholesterol metabolism, retinol metabolism (retinol, 18-hydroxyretinoic acid), vitamin and cofactor metabolism (thiamine, biotin, pantothenate, folate derivatives, riboflavin, nicotinamide, NAD/NADH, CoA), taurine and hypotaurine metabolism, propanoate metabolism, and branched-chain amino acid (BCAA) degradation (valine, leucine/isoleucine). Boxplots show scaled metabolite abundances; pairwise significance: *p < 0.05; **p < 0.01; ***p < 0.001; ****p < 0.0001; ns, not significant.

To distinguish which of these early metabolite alterations are microbiota-driven from those that are secondary to the emerging metabolic phenotype, we examined the same metabolites in the germ-free versus conventional comparison (Supplementary Figure 18). Most metabolites dysregulated in low-responders differed significantly by colonization status, including cholesteryl acetate, arachidonate, 18-hydroxyretinoic acid, the taurine/hypotaurine pair, branched-chain amino acids, and the microbial vitamin and cofactor derivatives thiamine, riboflavin, biotin amide, pantothenate, nicotinamide, and 5,10-methenyltetrahydrofolate, confirming that their hepatic abundance depends on microbial colonization, consistent with the loss of microbial biosynthetic capacity in the LiMa model. In contrast, a subset, including the NAD/NADH pool and propionic acid, did not differ between germ-free and conventional mice, suggesting that their early alteration may reflect host metabolic adaptation, potentially to caloric excess, rather than direct microbial provision.

## Discussion

Obesity is characterized by impaired metabolic plasticity, yet it remains unclear why some molecular alterations are reversed by lifestyle intervention whereas others persist, and how these persistent changes may contribute to weight regain. By integrating seven omic layers across the liver and gut in diet-induced obese mice, we identify two distinct axes of molecular remodeling in response to lifestyle intervention. The first is a reversible response dominated by host epigenetic and metallomic changes. The second is a persistent response that is strongly associated with loss of gut microbiota functional capacity. Together, these findings suggest that recovery from obesity involves both highly plastic host mechanisms and persistent alterations at the gut–liver interface.

Factor 1 captured the molecular changes most responsive to lifestyle intervention and was dominated by DNA methylation and histone modifications. Nearly 90% of HFD-induced DMRs returned toward control levels after intervention, highlighting the remarkable plasticity of the liver epigenome in response to dietary and behavioral changes. This contrasts with evidence from adipose tissue, where obesity-associated epigenetic alterations can persist after weight loss and contribute to an obesogenic memory^12^. These differences suggest that the reversibility of obesity-associated epigenetic remodeling may be tissue-dependent.

Notably, intervention did not simply restore HFD-induced hypermethylation to control levels; at a substantial number of loci, methylation was reduced below the levels observed in control animals. This pattern is consistent with previous reports of active DNA demethylation following exercise and caloric restriction in human skeletal muscle and adipose tissue^34,35^. The relevance of this response to human weight loss is further supported by a study identifying DNA methylation markers associated with the response to lifestyle-based weight loss in children with obesity^36^. Of the 214 responder-associated CpGs identified in that study, 32 genes overlapped with the reversible Factor 1 genes in our model, including *GFRA1* and *Gas7*, and showed comparable methylation patterns in mice and humans.

The coordinated recovery of histone modifications, including H3K27me3, a repressive mark previously associated with improved leptin sensitivity and reduced metabolic dysfunction^37^, together with the restoration of autophagy, apoptosis, and efferocytosis pathways, further supports extensive chromatin remodeling during metabolic recovery^38,39^. Importantly, approximately 15% of genes associated with gene-body DMRs encoded transcription factors, and their predicted regulatory networks encompassed ∼90% of the reversible response transcriptome. This suggests that methylation changes at a relatively small number of regulatory nodes could amplify the broad downstream transcriptional adaptation captured by Factor 1. Such hierarchical transcription factor architectures have been described in the liver, where a limited set of interconnected TFs controls broad transcriptional programs governing hepatocyte identity and metabolic function^40^. From a therapeutic standpoint, the identification of methylation-sensitive TF hubs governing the reversible response raises the possibility that targeted epigenetic editing at a small number of regulatory loci could recapitulate the transcriptional benefits of lifestyle intervention without requiring its full implementation^41,42^.

Our results also identify metals as regulators of metabolic plasticity, a layer that has received comparatively little attention in obesity research. Gut metal levels closely reflected dietary content, consistent with the lower micronutrient density of the high-fat diet. However, hepatic levels of Fe, Cu, Mn, Se, and Zn did not match dietary intake. Because the metal content of the high-fat and intervention diets was similar, the recovery of hepatic metal homeostasis following the lifestyle intervention cannot be explained by diet alone. Instead, it likely reflects active changes in metal absorption, hepatic storage, or systemic distribution that occur as metabolic health improves.

Selenium and manganese showed the largest number of negative associations with liver pathway variables within the reversible response, including efferocytosis, autophagy, and phagosome pathways, which are involved in inflammatory resolution and cellular stress adaptation^43,44^. Previous studies provide biological support for a role of these micronutrients in metabolic regulation. Selenium supplementation can protect HFD-fed mice against obesity, steatosis, and impaired glucose regulation^45^, while manganese supplementation^46^ has been associated with reduced hepatic lipid accumulation in animal models^47^ and with lower metabolic syndrome risk in men^48^. Although these associations do not establish a causal role for individual metals in the reversible response, they highlight the metallome as a potentially important and underexplored component of metabolic recovery.

In contrast to Factor 1, Factor 2 captured alterations that were resistant to lifestyle intervention. Two independent analyses point toward an early and predominantly microbiota-associated origin. First, germ-free filtering indicated that a substantial fraction of persistent hepatic changes falls within the microbiota-responsive space. Second, HFD-exposed low-responder mice showed that approximately half of the Factor 2 variables were already altered before the full obese phenotype developed. Together, these observations suggest that the persistent component is not simply a late molecular scar left by established obesity, but may represent an earlier feature of metabolic dysfunction associated with microbiota–liver interactions.

Remarkably, our study showed that taxonomic and functional recovery of the gut microbiota are uncoupled after lifestyle intervention. Although genus-level alpha diversity recovered, microbial functional capacity remained persistently reduced. This observation is consistent with the seminal work of Thaiss et al.^49^, which showed that weight loss can restore body weight and metabolic health without fully restoring microbiome gene content and function, suggesting a persistent functional memory of obesity. Consistent with this, microbial gene richness and functional capacity are increasingly recognized as more sensitive and clinically informative indicators of microbiome health than taxonomic diversity metrics, which are frequently insensitive to intervention^50,51^.

Our study suggests that the loss of numerically minor but functionally specialized taxa leaves a biosynthetic gap that dominant generalists cannot compensate. Moreover, the microbial functions depleted in our cohort mainly produce metabolites that the host cannot synthesize *de novo*. Humans and other mammals depend on dietary and microbial sources of several B vitamins, including cobalamin, folate, biotin, thiamine, and pantothenate^52,53^. These biosynthetic pathways are unevenly distributed across the microbiome. Cobalamin synthesis, for example, is energetically expensive and restricted to a limited number of taxa^54–57^, while many gut bacteria depend on B vitamins produced by these microbial specialists^58–61^. This metabolic interdependence may make the microbiome particularly vulnerable to the loss of specialized producer taxa. Consistent with this possibility, several genera depleted in our study, including Parasutterella^62,63^, Anaeroplasma^64^, Eisenbergiella^65^, Ruminococcus^66,67^, Marvinbryantia^68,69^, Acetatifactor^70^ are predicted B-vitamin producers. Their depletion could therefore reduce the biosynthetic capacity of the community even after broader taxonomic diversity has recovered.

B-vitamin deficiency has also been reported in obesity and type 2 diabetes, including biotin^71^, cobalamin^72,73^, and folate^74^. These abnormalities are often attributed to dietary factors or impaired absorption. Our findings raise an additional possibility: reduced microbial production may contribute to the persistent metabolic alterations associated with obesity. This possibility is particularly relevant because many of the hepatic pathways altered in Factor 2 depend on vitamin-derived cofactors. Peroxisomal β-oxidation, branched-chain amino acid degradation, steroid and cholesterol biosynthesis, retinol metabolism, and taurine metabolism require cofactors including CoA, FAD, NAD(P), or pyridoxal phosphate. Other processes, such as PPAR signaling and circadian regulation, may be affected indirectly through changes in lipid and redox metabolism. Thus, persistent loss of microbial biosynthetic capacity could potentially propagate to host metabolism by limiting the availability of microbial-derived vitamins or their downstream cofactors. We therefore propose a model in which loss of biosynthetically competent microbial taxa reduces microbial cofactor production and contributes to the persistence of hepatic metabolic alterations after obesity. This hypothesis remains to be tested experimentally, as our analyses establish association rather than causality.

Finally, cobalt provides an additional potential link between dietary composition, microbial metabolism, and the persistent response. Unlike Fe, Cu, Zn, Se, and Mn, which were associated primarily with the reversible component, cobalt was specifically associated with Factor 2 and remained low after lifestyle intervention. Cobalt remained low after the lifestyle intervention because the diet did not correct cobalt intake. This is particularly notable because cobalt has no essential metabolic role in mammals independent of cobalamin: it forms the metal center of vitamin B12, whose de novo synthesis is restricted to specific bacteria and archaea^57^. Host cobalt status may therefore be closely connected to microbial cobalamin metabolism.

The relevance of cobalt to metabolic health is supported by observations in humans and animal models. Higher circulating cobalt has been associated with lower insulin resistance in women with obesity^75^, while cobalt concentrations have been reported to be lower in women with obesity than in normal-weight controls^76^ and associated with obesity-related phenotypes in children^77^. In mice, cobalt administration has also been reported to attenuate lipid dysregulation and improve adipokine and glucose profiles following HFD exposure^78^.

The mechanism linking persistent cobalt deficiency to the Factor 2 phenotype remains unknown. One possibility is that sustained cobalt limitation could affect the abundance or activity of cobalamin-producing microorganisms, further constraining microbial B12 biosynthesis and thereby delaying functional recovery of the microbiome. In this model, persistent dietary cobalt deficiency could reinforce a pre-existing loss of microbial biosynthetic capacity, creating a feed-forward interaction between micronutrient availability, microbiota function, and hepatic metabolism. This hypothesis remains to be tested, but it provides a potential mechanistic framework linking dietary micronutrients to the persistence of obesity-associated metabolic alterations.

## Author Contributions

J.R., J.M-B and O.Y. designed the study. J.M-B performed experiments, collected and pre-processed the data. J.R and O.Y. designed the data analysis strategy. J.R. analyzed and integrated all the data. J.R. and O.Y. wrote the paper. A.B, C.M, and S.R conducted the *in vivo* experiment and tissue collection for the germ-free mouse model. P.G-P and P.M.G-R conducted the *in vivo* experiment and tissue collection for the LiMa model. B.C and P.G-T conducted the shotgun metagenomics experiments. J.C. supervised and analyzed metabolomics data. I.F and A.I supervised the histone proteomics experiments. C.M.H and M.S supervised and analyzed the DNA methylation experiments. L.S and M.M supervised and analyzed the RNAseq experiments

## Acknowledgments

This work was supported by the European Union’s Horizon 2020 research and innovation program under the Marie Skłodowska-Curie grant agreement No 675610 (ChroMe, MSCA-ITN-2015) and No. 824110 (EASI-Genomics). O.Y. acknowledges the financial support of grants BFU2017-87958-P and PID2022-136226OB-I00 funded by MICIU/AEI/10.13039/501100011033 and by the European Union NextGenerationEU/PRTR. J.R. acknowledges the financial support of the Government of Catalonia through the predoctoral grant 2023FI-2 00620. P.M.G.-R acknowledges the financial support of grants PI15/00701 from Instituto de Salud Carlos III (ISCIII) and PID2022-138537OB-I00 from Ministerio de Ciencia e Innovación (MICINN); cofinanced by the European Regional Development Fund ‘‘A way to build Europe’’. The Government of Catalonia supports AGAUR 2017-SGR-204 and 2021SGR842 grants.

## Methods

### Lifestyle Matters (LIMA) model

A subset of 36 mice was selected from the Lifestyle Matters (LiMa) project, comprising 10 animals from each of the three main experimental groups, control (CTR), high-fat diet (HFD), and intervention (INT), and 6 animals from the high-fat diet-excluded group (HFDex). The LiMa experimental design has been described in detail elsewhere^25^ and is summarized here. Male C57BL/6JOlaHSD mice (Envigo, IN, USA) were housed under a 12 h light/12 h dark cycle. Mice were initially divided into two dietary groups for 16 weeks: a control group fed a standard chow diet (Teklad Global 14% Protein Rodent Maintenance Diet, Envigo), and a high-fat diet group (HFD; D12451, Research Diets, NJ, USA). Inclusion criteria for the CTR group included body weight (BW) < 33 g, normoglycemia, normal glucose tolerance, insulin sensitivity, and fasting insulin levels. Mice fed the high-fat diet (HFD) were required to meet predefined criteria of metabolic impairment, including body weight >37 g, fasting hyperglycemia, hyperinsulinemia, insulin resistance, and glucose intolerance. Animals that did not meet these criteria were classified as the high-fat diet excluded (HFDex) group. A subset of the metabolically impaired HFD mice was then subjected to a lifestyle intervention consisting of dietary modification and structured exercise. The diet was switched to a modified formulation (Research Diets, NJ, USA) maintaining 45% caloric intake from fat, with replacement of simple carbohydrates (sucrose) by complex carbohydrates (corn starch) and animal-derived fats (lard) by plant-based oils (olive oil and flaxseed), increasing mono- and poly-unsaturated fatty acids, including omega-3. The intervention was conducted in two phases. The first phase (5 weeks) included caloric restriction (80% of CTR intake during the first week, followed by 100% for the remaining 4 weeks) combined with treadmill exercise (Exer-6M, Columbus Instruments, OH, USA), performed 5 days/week, for 1 h/day. Exercise intensity was progressively increased up to 20 m/min with a 10° incline. The second phase (additional 5 weeks) maintained the dietary intervention and implemented an adjusted exercise regimen consisting of alternate-day sessions (1 h/day) at 16 m/min and 5° incline. Mice from this second phase were selected as the intervention group (INT) in this study. All experimental procedures were approved by the Comitè Ètic d’Experimentació Animal of the Universitat de Barcelona and the Departament d’Agricultura, Ramaderia, Pesca, Alimentació i Medi Natural of the Generalitat de Catalunya, and were conducted in accordance with European and Spanish regulations.

### Germ-Free mouse model

nine-week-old male germ-free (GF) C57BL/6J mice (n=10) were obtained from Anaxem (Micalis Institute, INRAE, Jouy-en-Josas, France; license B78-33-6) and housed in sterile isolators (Getinge, Les Ulis, France). Conventionally raised (CV) C57BL/6J control mice (n=10) were purchased from Charles River Laboratories (L’Arbresle, France), transported to Anaxem, and maintained under comparable conditions in non-sterile isolators to control for environmental factors. Mice were housed at 20–24°C under a 12 h light/dark cycle with ad libitum access to autoclaved water and gamma-irradiated chow diet (45 kGy; R03, Scientific Animal Food and Engineering, Augy, France). Germ-free status was verified weekly by microscopy and culture-based screening of fecal samples. Experiments were conducted in two randomized batches to minimize confounding effects. All procedures were approved by the ethics committee of the INRAE Research Center at Jouy-en-Josas and complied with European regulations

### Liver sample preparation

liver tissues were collected, immediately snap-frozen in liquid nitrogen, and stored at −80°C. Samples were subsequently lyophilized, pulverized, and homogenized prior to nucleic acid and metabolite extraction.

### Transcriptomics (RNA sequencing)

total RNA was extracted from ∼12 mg of lyophilized liver tissue using the Direct-zol RNA Miniprep Plus kit (Zymo Research, CA, USA), including on-column DNase I treatment. Briefly, samples were homogenized in TRI reagent, followed by ethanol addition and column-based purification according to the manufacturer’s protocol. RNA was eluted in RNase-free water. RNA quantity and quality were assessed using NanoDrop 2000 (Thermo Fisher Scientific), Qubit RNA HS/BR assays (Thermo Fisher Scientific), and Agilent Bioanalyzer 2100 (RNA 6000 Nano kit), ensuring high RNA integrity (RIN).

### Library preparation and sequencing

ribosomal RNA was depleted using the Illumina Ribo-Zero Plus kit. Strand-specific libraries were prepared with the TruSeq Stranded Total RNA kit (Illumina) following the manufacturer’s instructions, including cDNA synthesis, adapter ligation, and PCR amplification. Libraries were validated using the Agilent Bioanalyzer (DNA 7500 kit) and sequenced on an Illumina HiSeq 4000 platform (paired-end, 75 bp), generating >60 million reads per sample. Base calling was performed using Real Time Analysis (RTA v2.7.7), and FASTQ files were generated for downstream analysis.

### RNA-seq data processing

sequencing quality was assessed using FASTQC (v0.11.5) and MultiQC (v1.8). Reads were aligned to the mouse reference genome (GRCm38) using STAR (v2.7.0f) with default parameters. Alignment quality was evaluated with QualiMap (v2.2.1). Gene-level counts were obtained using HTSeq (v0.11.2).

### Differential gene expression analysis

raw count data were imported into R (v4.2.2) and filtered to retain transcripts detected (>0 counts) in at least 80% of samples within any group and with expression >1 TPM. Differential expression analysis was performed using DESeq2 (v1.40.0). For the LiMa dataset, a ∼0 + group design was used, whereas the germ-free dataset included batch as a covariate (∼0 + group + batch). Wald tests were conducted for the following contrasts: HFD vs CTR, INT vs CTR, and INT vs HFD (LiMa), and GF vs CV (germ-free model). False discovery rate (FDR)-adjusted p-values were reported.

### Histone modifications proteomics

#### Sample preparation

histone modifications were analyzed from ∼14 mg of lyophilized and homogenized liver tissue per sample. Histones were acid-extracted, digested, desalted, and analyzed by LC–MS following a previously described protocol^79^, with modifications to the initial extraction steps. Briefly, samples were resuspended in 150 µL of 0.2 M H_2_SO_4_, sonicated (Bioruptor, Diagenode; 15 cycles of 30 s on/off, high setting), and incubated at 4°C under rotation. After centrifugation (30 min, 14,000 rpm, 4°C), supernatants were processed according to the referenced protocol to precipitate, wash, and resolubilize histones, followed by separation using a polyacrylamide gel electrophoresis. For protein digestion, gel bands corresponding to histones were excised and subjected to in-gel acylation and trypsin digestion. Peptides were extracted using formic acid to maximize recovery. Prior to LC–MS analysis, peptides were desalted using sequential C18 StageTips and carbon columns with intermediate washing steps.

#### LC–MS analysis and data processing

peptides were analyzed by liquid chromatography–mass spectrometry using a C18 HPLC column coupled to a Q Exactive HF Orbitrap instrument (Thermo Fisher Scientific, MA, USA) operated in parallel reaction monitoring (PRM) mode, alternating between MS1 and MS2 scans. No isotopically labeled standards were used. Raw data were processed using Skyline (v4.2.0.19072). MS1 and MS2 signals were matched to a spectral library based on retention time and transition values and manually validated using GPMAW (v5.02) and Xcalibur Qual Browser (v4.1.31.9) (Thermo Fisher Scientific, MA, USA). Peptide identifications required at least four matching MS2 fragment ions. Relative abundances of histone modifications were calculated using the “peptide family” approach, whereby MS1 peak areas for peptides sharing the same sequence backbone were normalized to the total signal of their corresponding peptide family.

#### Statistical analysis

differences in histone modification levels across experimental groups were assessed using the Kruskal–Wallis test. When significant (p < 0.05), pairwise comparisons were performed using the Conover post hoc test (DescTools v0.99.54), with p-values adjusted for multiple testing using the false discovery rate (FDR) method.

### DNA Methylation

#### DNA extraction and quality control

genomic DNA was extracted from liver tissue using the QIAamp DNA Mini Kit (Qiagen, Hilden, Germany). DNA quality was assessed using the Agilent 2100 Bioanalyzer (DNA 12000 kit), and concentration was measured with the Qubit dsDNA assay on a Qubit 4.0 fluorometer (Thermo Fisher Scientific, MA, USA).

#### Library preparation and sequencing

whole-genome bisulfite sequencing (WGBS) libraries were prepared using the Accel-NGS Methyl-Seq DNA library kit (IDT, Coralville, IA, USA) following the manufacturer’s instructions. Libraries were sequenced on an Illumina NovaSeq 6000 platform (paired-end, 100 bp reads) (Illumina, San Diego, US).

#### Data processing

raw sequencing data were processed using the nf-core/methylseq pipeline (v2.5.0)^80–82^. Briefly, read quality was assessed with FastQC (v0.11.9), and adapter trimming was performed using Trim Galore! (v0.6.7). Reads were aligned to the mouse reference genome (mm39) using Bismark (v0.24.0), followed by deduplication and extraction of methylation calls. Coverage files generated by Bismark, were imported into R using the methrix package (v1.16.0), applying the *Bismark_cov* workflow with stranded = TRUE, collapse.strands = TRUE, and zero.based = FALSE. CpG sites were annotated using the BSgenome.Mmusculus.UCSC.mm39 reference (v1.4.3). Low-coverage CpGs (<5 reads) were masked, and uncovered CpGs were removed. Samples with abnormally low coverage were excluded from downstream analyses.

#### Differential methylation analysis

differentially methylated regions (DMRs) were identified using the DSS package. A multi-factor model was fitted at the CpG level to assess differences across experimental contrasts: HFD vs CTR, INT vs CTR, and INT vs HFD (LiMa model), and GF vs CV (germ-free model). For each comparison, DMRs were defined as genomic regions containing ≥50% significant CpGs (unadjusted p < 0.001). To enable cross-condition comparisons, DMRs from all contrasts were merged into non-overlapping regions. Mean methylation levels were then calculated for each region across samples. Genomic annotation of DMRs was performed using ChIPseeker (v1.38.0) with the TxDb.Mmusculus.UCSC.mm39.knownGene database (v3.18).

### Shotgun metagenomics

#### DNA extraction and sequencing

genomic DNA from colon and cecum contents was extracted using the MagMAX CORE Nucleic Acid Purification Kit (Thermo Fisher Scientific, CA, USA). Sequencing libraries were prepared from 1 ng of DNA using a Tn5 transposase-based fragmentation approach (Nextera workflow). Adapter-ligated fragments were PCR-amplified using KAPA HiFi HotStart ReadyMix (Kapa Biosystems) with dual indexing (i5 and i7 barcodes), followed by purification with AMPure XP beads (Beckman Coulter). Library quality and fragment size distribution were assessed using the Agilent Bioanalyzer and Fragment Analyzer (High Sensitivity assay), and quantified by qPCR (KAPA Library Quantification Kit). Libraries were sequenced on an Illumina HiSeq 2500 platform to generate 2 × 150 bp paired-end reads.

#### Sequencing data processing

raw read quality was assessed using FastQC (v0.11.8), and adapter trimming and filtering were performed with Trimmomatic (v0.39). For taxonomic profiling, 16S rRNA sequences were extracted using SortMeRNA (v4.2.0) with SILVA reference databases (Archaea and Bacteria), and taxonomy was assigned using MAPseq (v2.0.1) with the SILVA v132 database. For functional analysis, reads were merged using BBMerge (v38.84), quality-filtered with Trimmomatic, and assembled de novo using MEGAHIT (v1.2.9). Gene prediction and annotation were performed with Prokka (v1.14.6), which uses Prodigal for open reading frame detection and multiple databases (including ISfinder, NCBI AMR, and UniProtKB/Swiss-Prot) for functional annotation. Clusters of Orthologous Groups (COGs)^83^ were assigned to coding sequences, and gene abundances were quantified by mapping reads back to assembled contigs using Bowtie2 (v2.3.5).

#### Microbiome data analysis

taxonomic and functional profiles were imported into R for downstream analysis. Features (genera and COGs) with low prevalence (<80% presence within any experimental group) and unclassified taxa were removed. Alpha diversity metrics (Shannon entropy and observed richness) were calculated using the *mia* package (v1.10.0) and compared using the Wilcoxon test. Beta diversity was assessed using principal coordinates analysis (PCoA) based on Aitchison distances, appropriate for compositional data, and statistical significance was evaluated using PERMANOVA (adonis2 function, vegan v2.6-4). Differential abundance analysis at the genus and functional (COG) levels was performed using DESeq2, applying the same model design and contrasts as in the RNA-seq analysis. Wald tests were used for significance testing, and p-values were adjusted for multiple comparisons using the false discovery rate (FDR) method.

### Metallomics (ICP-MS)

#### Sample preparation

liver and gut samples were prepared by cryogenic homogenization with a SPEXSamplePrep (Freezer, 6770) cryogenic homogenizer for 30 s at 10 strokes per second and were mineralized in a microwave oven. In summary, 0.1 g of each tissue was put into MiniXpress polytetrafluoroethylene (PTFE) 5 mL tubes, 0.5 mg of a nitric acid and hydrogen peroxide mixture (4:1; v/v) were added, 1 µg/L of Rhodium was added to use as an internal standard, samples were pre-digested for 10 min and mineralized in a MARS microwave oven (CEM, Matthews, USA) with a 15 min temperature ramp from room temperature to 160°C that was held for 20 min.

#### ICP-MS analysis

metal samples absolute concentrations were then quantified by Inductively Coupled Plasma Mass Spectrometry (ICP-MS) model 8800 Triple Quad ICP-MS (Agilent Technologies, Tokyo, Japan). Skimmer cones and nickel sampling were utilized with a sampling depth of 10 mm, forward power was set to 1550 W; gas flow rates were 15 L/min for plasma gas and 1.08 L/min for carrier gas; a solution of 1 µg/L Li, Co, Y and TI was utilized for calibration; and high purity helium (He), oxygen (O_2_) (>99.999%) and hydrogen (H_2_) (>95%) were employed to eliminate interferences. Helium flow rate was 4.5 mL/min for most elements, except for selenium, that required a mix of H_2_ (2 mL/min) and O_2_ (40%) in MS/MS. The following isotope elements were monitored with a dwell time of 0.3 s/isotope: ^24^Mg, ^27^Al, ^51^V, ^52^Cr, ^55^Mn, ^56^Fe, ^59^Co, ^63^Cu, ^66^Zn, ^75^As, ^78^Se, ^80^Se, ^95^Mo, ^103^Rh, ^111^Cd, and ^208^Pb.

#### Statistical analysis

absolute abundances of each metal were compared across levels with non-parametric tests: a Kruskal-Wallis test between all groups, followed with a Connover test (DescTools v0.99.54) for pairwise comparisons and p-values adjusted with an FDR.

### Untargeted metabolomics (LiMa and germ-free cohorts)

#### Sample preparation

approximately 7.5 mg of homogenized liver tissue was extracted with 400 µL of acetonitrile:methanol (4:4:2, v/v/v). Samples were vortexed for 1 min and subjected to three extraction cycles consisting of liquid nitrogen incubation (30 s), water-bath sonication (30 s), and vortex mixing (30 s). Extracts were incubated at −20 °C for 60 min and centrifuged at 22,000 × g for 10 min at 4 °C. Subsequently, 100 µL of supernatant was transferred to LC–MS vials for analysis.

#### LC–MS analysis

metabolomic profiling was performed on an Agilent 1200 Series UHPLC system coupled to an Agilent G6550A ESI-qTOF mass spectrometer operating in positive and negative electrospray ionization modes. Metabolites were analyzed using both hydrophilic interaction chromatography (InfinityLab Poroshell 120 HILIC-Z, 2.1 × 100 mm, 2.7 µm, Agilent Technologies) and reversed-phase chromatography (ACQUITY UPLC HSS T3, 2.1 × 150 mm, 1.8 µm, Waters). For HILIC separations, mobile phase A consisted of water containing 50 mM ammonium acetate and mobile phase B consisted of acetonitrile. The gradient was as follows: 98% B for 0–2 min, decreased to 40% B from 2–10 min, returned to 98% B between 10–10.5 min, and maintained at 98% B until 15 min. For reversed-phase chromatography, mobile phase A consisted of water containing 0.1% formic acid. The gradient increased to 100% A over 0–7.5 min, was held from 7.5–8.5 min, and returned to the initial conditions between 8.5–10 min. Mass spectrometry was performed with the following source parameters: gas temperature, 200 °C; drying gas, 14 L min−1; nebulizer pressure, 35 psig; fragmentor voltage, 175 V; and skimmer voltage, 65 V. Data were acquired from *m/z* 50–1100 at 3 spectra s−1. Quality control (QC) samples, prepared by pooling aliquots from all study samples, were injected throughout the analytical sequence to monitor instrument performance and correct for signal drift.

#### Data processing

raw data files were converted to mzML format using msConvert (ProteoWizard v3.0.21034). The three analytical datasets (HILIC positive, HILIC negative and reversed-phase positive) were processed independently using the same workflow with minor parameter adjustments. Peak detection and alignment were performed in R using XCMS (v4.0.2). Chromatographic peaks were detected with the CentWave algorithm (10 ppm mass accuracy; peak width 5–50 s), refined using MergeNeighboringParams, aligned with the ObiWarp algorithm, grouped into chromatographic features (minFraction = 0.4, binwidth = 5, binsize = 0.01), and missing peaks were recovered using fillChromPeaks. Features were subsequently filtered using the pmp package (v1.14.1). Only features detected in at least 80% of biological samples and 30% of QC samples, showing at least a two-fold difference between samples and blanks (log2 fold change ≥1), and exhibiting a QC relative standard deviation (RSD) ≤35% were retained. Signal drift across the analytical sequence was corrected using the QCRSC algorithm implemented in the pmp package (v1.14.1).

#### Statistical analysis

for the LiMa cohort, differences among experimental groups were evaluated using the Kruskal–Wallis test. Significant features (*P* < 0.05) were subsequently analyzed by pairwise Conover tests, and *P* values were adjusted for multiple testing using the Benjamini–Hochberg false discovery rate (FDR). For the Germ-Free dataset, differential analysis was performed using an ANOVA model including batch and experimental group as fixed effects (∼ batch + group). Pairwise comparisons were performed using Tukey’s honestly significant difference test, and *P* values were corrected using the Benjamini–Hochberg FDR procedure. Model residuals were inspected to verify approximate homoscedasticity and normality.

#### Feature annotation

metabolite annotation was performed using RHermes (v0.99.0)^84^. A reference database combining HMDB, ChEBI, di- and tripeptide database was used to generate a list of theoretical adducts with setDB. The adduct list included M+H, M+NH4, M+Na, M+K and M+ for positive ionization, and M−H and M+Cl for negative ionization. Three pooled QC samples and one blank sample were processed with processMS1 using a mass tolerance of 10 ppm, 30,000 resolving power, *m/z* range 50–1500, qTOF instrument settings and a minimum intensity threshold of 1,000. Scans of interest (SOIs) were generated using the double-detection mode, and blank samples were used to remove background signals. SOIs with isotopic fidelity ≥0.8 and intensity ≥1,000 were retained. Features detected in at least two of the three QC samples were merged into a consensus SOI list and matched to XCMS features using qHermes (v0.0.0.90). Complementary annotation was performed with mummichog (Python v3.8.13), using default parameters except for ionization mode. Feature ranking was based on Kruskal– Wallis statistics for the LiMa dataset and ANOVA *t*-statistics for the Germ-Free dataset. Annotations obtained with RHermes and mummichog were combined to identify redundant signals. Confirmed isotopes lacking independent annotations and multiple adducts assigned to the same metabolite with overlapping retention times were collapsed into a single representative feature. Whenever possible, the protonated ion (M+H or M−H) was retained. If no protonated ion was detected, the adduct showing the strongest statistical association (ANOVA for the Germ-Free cohort or Kruskal–Wallis for the LiMa cohort) was selected as the representative feature.

### Targeted Metabolomics

#### LC–MS analysis

Targeted metabolites were analyzed using an Agilent InfinityLab Poroshell 120 HILIC-Z column (2.1 × 100 mm, 2.7 μm; Agilent Technologies) maintained at 25 °C and coupled to an Agilent 6490 triple quadrupole (QqQ) mass spectrometer operating in simultaneous positive and negative electrospray ionization modes. Mobile phase A consisted of water containing 50 mM ammonium acetate and medronic acid, whereas mobile phase B consisted of acetonitrile. The flow rate was 0.4 mL min−1 with the following gradient: 98% B (0–2 min), decreased to 40% B (2–9 min), returned to 98% B (9–9.5 min), and held until 13 min for column re-equilibration. Mass spectrometry was performed using the following source parameters: nebulizer pressure, 35 psi; drying gas temperature, 270 °C; sheath gas temperature, 400 °C; drying gas flow, 15 L min−1; sheath gas flow, 11 L min−1; capillary voltage, ±3,000 V; nozzle voltage, +1,000 V (positive mode) and −1,500 V (negative mode). The iFunnel settings were HRF/LRF 130/100 V in positive mode and 110/60 V in negative mode. A 5 μL injection volume was used. Monitored precursor/product ion transitions and collision energies are listed in Supplementary Table 8. Pooled quality control (QC) samples were injected throughout the analytical sequence to monitor analytical performance and correct for signal drift. Chromatographic peaks were manually integrated using MassHunter Qualitative Analysis (Agilent Technologies)^85^.

#### Liver short-chain fatty acid (SCFA) extraction and quantification

25 mg of liver tissue was transferred to 1.5 mL Eppendorf tubes. To each sample, 60 µL of internal standard (butyric acid-LAB, acetic acid-LAB, and propionic acid-LAB in methanol), 333 µL of water:methanol (1:1, v/v), 666 µL of chloroform, 100 µL of methanol, and a steel bead were added. Samples were homogenized in a bullet blender (1 min, speed 7), vortexed for 1 min, and centrifuged (5 min, 15,000 rpm, 4 °C). A derivatization step was then performed: 40 µL of supernatant was combined with 40 µL of water, 10 µL of 0.1 M BHA, and 10 µL of 0.25 M EDC, and incubated at 25 °C for 1 h. Following incubation, 600 µL of diethyl ether was added, and samples were vortexed for 10 min and centrifuged (5 min, 15,000 rpm, 4 °C). The upper organic layer (400 µL) was transferred to a new tube, evaporated under a nitrogen stream, reconstituted in 200 µL of water:methanol (1:1, v/v), and transferred to vials for HPLC analysis. Chromatographic separation was performed at 45 °C on a reverse-phase Kinetex 2.6 µm polar C18 column (100 Å, 100 × 2.1 mm; Phenomenex, Torrance, CA, USA) fitted with a guard precolumn. Mobile phase A consisted of 0.1% formic acid and 10 mM ammonium formate in water; mobile phase B consisted of 0.1% formic acid in methanol:isopropanol (9:1, v/v). At a flow rate of 0.3 mL/min, the gradient was: isocratic 32–60% B (0–4.6 min), increase to 65% B (4.6–5.5 min), increase to 98% B (5.5–7 min), held at 98% B (7–9 min), and return to 32% B (9– 10 min), held for 1 min to re-equilibrate the column. Chromatographic peaks were integrated and quantified automatically using the Qualitative Analysis module of MassHunter Workstation (Agilent Technologies), and absolute concentrations were calculated using the labeled internal standards.

#### Data processing and statistical analysis

Peak areas were imported into R and corrected for injection-order drift using the QCRSC algorithm implemented in the pmp package (v1.14.1). Signal intensities were normalized to the exact liver mass used for extraction. Multivariate outliers were identified by Hotelling’s T² 99% confidence ellipse and removed prior to statistical analysis. Group differences were assessed using the Kruskal–Wallis test followed by pairwise Conover tests for significant metabolites (*P* ≤ 0.05), with false discovery rate (FDR) correction for multiple testing.

### Multi-Omics Factor Analysis (MOFA)

#### Omic layer pre-processing

data from all omics were integrated into MultiAssayExperiment, one for the GF model data and another for the LiMa. Each omic layer was processed with the following ordered steps: imputed using a KNN with 10 neighbors approach for all the omic layers that contained missing values; normalized using a variance stabilizing transformations (vst) from the DeSeq package (v1.40.0) only for the RNA-Sequencing, Untargeted Metabolomics and Shotgun metagenomics data; batch corrected with ComBat (sva v3.50.0) for the GF mice Untargeted Metabolomics and RNA-Sequencing layers; and all layers were scaled and centered to ensure comparability between variables.

### MOFA models

#### Intervention (INT) model training

All LiMa omic layers were filtered to retain variables that were statistically significant in at least one group comparison (HFD–CNT, HFD–INT, or INT–CNT), and multi-omics factor analysis models were built using the MOFA2 package (v1.12.1). Models were trained with default parameters, except that the convergence mode was set to "slow" and the maximum number of iterations to 5,000. To determine the appropriate number of factors, models were trained with increasing numbers of factors (2, 3, 4, 5, and 10) to assess factor consistency, together with an additional model restricted to the three largest omic layers (untargeted metabolomics, RNA sequencing, and DNA methylation) to check for sample-size bias. For each configuration, three models were trained with different starting seeds to avoid convergence to local minima, and the model with the lowest ELBO score was retained. After comparing factor scores across all trained models (Supplementary Figure 2 and Figure 1), the two-factor model was selected, as additional factors did not resolve any distinct data patterns.

#### HFDex model training

For the second MOFA model, the INT group samples were replaced with the HFDex samples, the same variables selected in the first model were used as input, and the model was trained across an increasing number of factors exactly as described above. As in the first model, the two-factor solution was selected, since additional factors did not uncover distinct data patterns.

#### Variable selection

Variables were assigned to factors using two criteria derived from their MOFA weights. First, each variable was classified by the sign concordance of its Factor 1 and Factor 2 weights: concordant (both positive or both negative) or non-concordant (opposite signs). Second, a factor-belonging score was computed as the absolute Factor 1 weight divided by the sum of the absolute Factor 1 and Factor 2 weights, yielding a value from 0 (fully Factor 2) to 1 (fully Factor 1). For each concordance class, the predicted mean variable intensity was plotted across the belonging score in 0.1 increments (0 to 1; Supplementary Figure 3), and variables were assigned by visual inspection of these thresholds: variables with a belonging score of 0.6–1 (both concordant and non-concordant) were assigned to Factor 1; concordant variables with a score of 0–0.3 and non-concordant variables with a score of 0–0.4 were assigned to Factor 2; and all remaining variables were assigned to neither factor. For the HFDex MOFA model, the variable-selection procedure applied to the first model was used, with model-specific thresholds (Supplementary Figure 16): non-concordant variables with a belonging score of 0.6–1 were assigned to Factor 1 and 0–0.4 to Factor 2; concordant variables with a score of 0.5–1 were assigned to Factor 1 and 0–0.1 to Factor 2.

#### Functional enrichment analysis

Gene Ontology (GO) enrichment of Entrez gene IDs was performed using the enrichGO function from clusterProfiler (v4.10.1), with the org.Mm.eg.db database (v3.18.0), all three ontologies (MF, BP, CC), a p-value cutoff of 0.05, FDR correction, a maximum gene-set size of 500, and all database genes as the background universe. Redundant GO terms with a semantic similarity greater than 0.7 were then collapsed, retaining only the most significant term in each group, and log2 enrichment values were calculated. KEGG gene-level and joint gene–metabolite pathway enrichments were computed analogously using the enricher function from clusterProfiler, with the enrichment database replaced by KEGG gene and KEGG gene– metabolite association collections, respectively, obtained via the KEGGREST package (v1.42.0). These analyses used the annotated putative KEGG metabolite IDs (excluding features without an assigned KEGG ID) together with gene Entrez IDs.

#### Correlation analysis

Pearson correlations between omic layers were calculated using the getExperimentCrossAssociation function from the mia package (v1.10.0), corrected for multiple testing (FDR < 0.05). To simplify visualization, the strongest significant correlations were retained (Pearson |r| ≥ 0.75–0.8) and displayed as chord diagrams (circlize, v0.4.16) and Sankey diagrams (networkD3, v0.4). Full correlation data are provided in Supplementary Files 6-7, 10, 14-15, 18.

#### Permutation tests

##### Directional bias of DMR–gene correlations

To test whether gene-body DMRs were preferentially associated with negative correlations, we generated an empirical null distribution by randomly sampling 10,000 sets of correlations from all significant correlations, with each set matched to the number of gene-body DMR–gene correlations observed. For each random set, we calculated the percentage of negative correlations. The observed percentage was then compared with the empirical null distribution to derive an empirical p-value. This analysis was performed separately for each genomic region (promoter, intron, exon, 5′ UTR, 3′ UTR, and distal intergenic regions within 1 Mb).

##### TF enrichment

To test whether the proportion of transcription factors (TFs) among the genes associated with gene-body DMRs or Factor-associated genes differed from that expected by chance, we first defined the list of TFs from the gene list by querying the TFCheckpoint 2.0 database, then we generated empirical null distributions by randomly sampling 10,000 gene sets from the corresponding background distribution. Depending on the analysis, the background comprised either all expressed genes (n = 11,797) or the Factor-associated genes. Each random set was matched to the size of the corresponding observed gene set, and the percentage of TFs was calculated for each permutation. The observed percentage was then compared with the empirical null distribution to derive an empirical p-value.

##### Overlap between variable sets

To test whether the overlap between two variable sets exceeded that expected by chance, we generated an empirical null distribution by independently sampling 10,000 pairs of variable sets from the full dataset, with each set matched to the size of the corresponding observed set. The number of overlapping variables was calculated for each permutation, and the observed overlap was compared with the resulting empirical null distribution to derive an empirical p-value. For example, to assess the overlap between GF–CV significant genes (n = 3,198) and INT Factor 2 genes (n = 1,913), we independently sampled 3,198 and 1,913 genes from all measured genes in each permutation and calculated their overlap. The resulting null distribution (Supplementary Figure 11C) was used to evaluate the observed overlap of 627 genes. The same procedure was applied to the overlap between GF–CV and LiMa metabolite features (Supplementary Figure 11B,C) and to comparisons between factors across models, with the corresponding sample sizes and null distributions shown in Supplementary Figure 16.

#### TFLink analysis

The TFLink *Mus musculus* small- and large-scale interaction table (version 1.0) was downloaded from the official TFLink website. Entrez gene IDs were mapped to Ensembl gene IDs using the mapIds function from the AnnotationDbi package (v1.64.1). Interactions were then filtered to retain high-confidence, experimentally supported interactions detected by chromatin immunoprecipitation. The resulting interactions were represented as a directed *igraph* network, with edges directed from transcription factors (TFs) to their target genes (TF → gene). The network was then matched to the Factor 1 dataset, and nodes not detected in the Factor 1 dataset were excluded.

##### TFLink network coverage

To quantify the proportion of Factor 1 genes covered by the TFLink network at distance 1, the shortest-path distance from each selected TF to all other nodes in the network was calculated using *igraph*. For each gene, the minimum non-zero distance from any selected TF was determined. Network coverage at distance 1 was defined as the number of genes with a minimum distance of 1 divided by the total number of genes in the network.

##### TFLink coverage permutation test

To assess whether the observed network coverage exceeded that expected by chance, we generated an empirical null distribution by randomly sampling 22 TFs (the number of TFs associated with gene-body DMRs matched to the TFLink network) from the 348 TFs represented in the network. This sampling procedure was repeated 10,000 times, and network coverage at distance 1 was calculated for each random TF set. The observed coverage obtained from the 22 gene-body DMR-associated TFs was then compared with the empirical null distribution to derive an empirical p-value (Supplementary Figure 8).

