## Supplementary Figures for "Multi-omic dissection of reversible and persistent molecular alterations in diet-induced obesity"

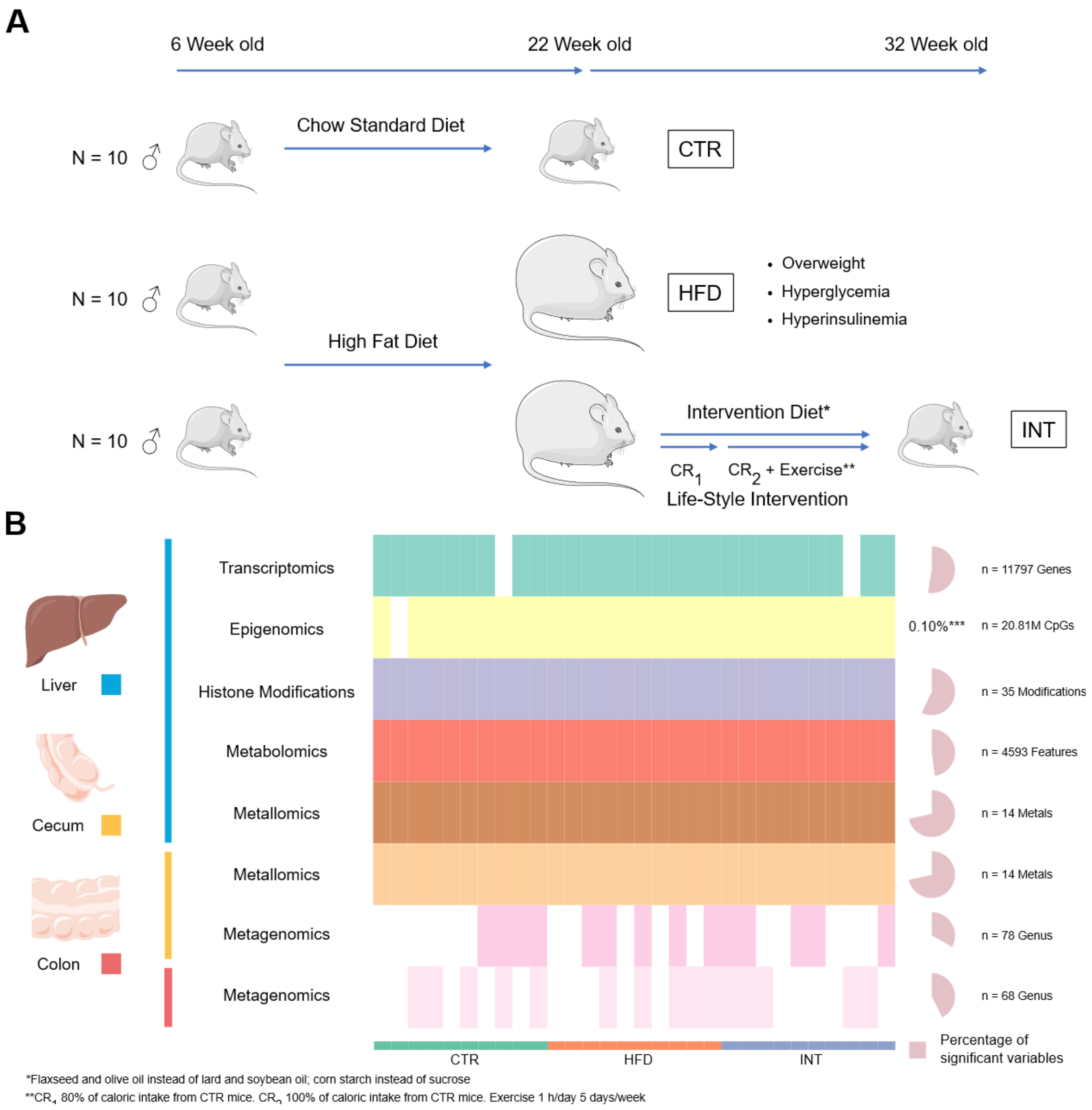


**Supplementary Figure 1**. **A. Experimental design of the Life-Style Matters (LiMa) model.** Six-week-old C57BL/6 mice were fed ad libitum either a standard chow diet (CTR) or a high-fat diet (HFD) for 16 weeks. A subset of HFD mice then underwent a 10-week lifestyle intervention (INT), consisting of dietary modification (replacement of lard with flaxseed and olive oil, and sucrose with corn starch), caloric restriction (80% of CTR intake during the first week, gradually increased for 4 weeks until 100% CTR intake, and last 5 weeks matching CTR intake), and exercise (first 4 weeks, 5 days/week for 1 h/day with exercise intensity progressively increased up to 20 m/min and a 10° incline, and the last 5 weeks an exercise regimen consisting of alternate-day sessions (1 h/day) at 16 m/min and 5% incline 5 days/week). **B. Overview of the collected multi-omics data.** From top to bottom: liver transcriptomics, epigenomics (DNA methylation), histone modifications, metabolomics, and metallomics; cecum metallomics and metagenomics; and colon metagenomics. Each square indicates sample availability for a given omic layer, and the lower bar denotes experimental groups (CTR, green; HFD, red; INT, blue). Numbers on the right indicate the total number of variables per omic layer, and the pie chart shows the proportion of variables significantly altered in at least one group comparison. ***DNA methylation changes correspond to 20,426 CpGs (from 4,591 DMRs) out of 20.81 million genome CpGs.

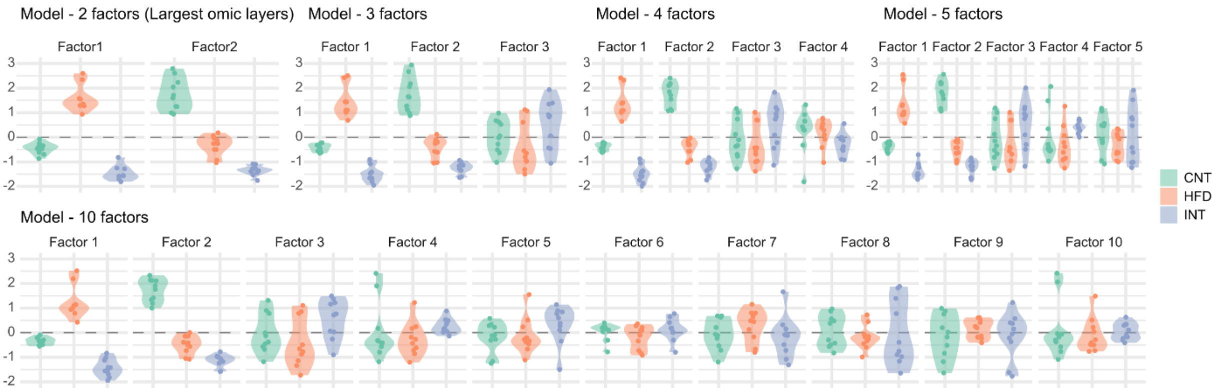


**Supplementary Figure 2. A**. Factor scores boxplots of the MOFA model trained with only the omics with the largest variables and 2 factors (untargeted metabolomics, DNA methylation and RNA sequencing) and an increasing number of factors divided by experimental groups (3, 4, 5 and 10 factors respectively).

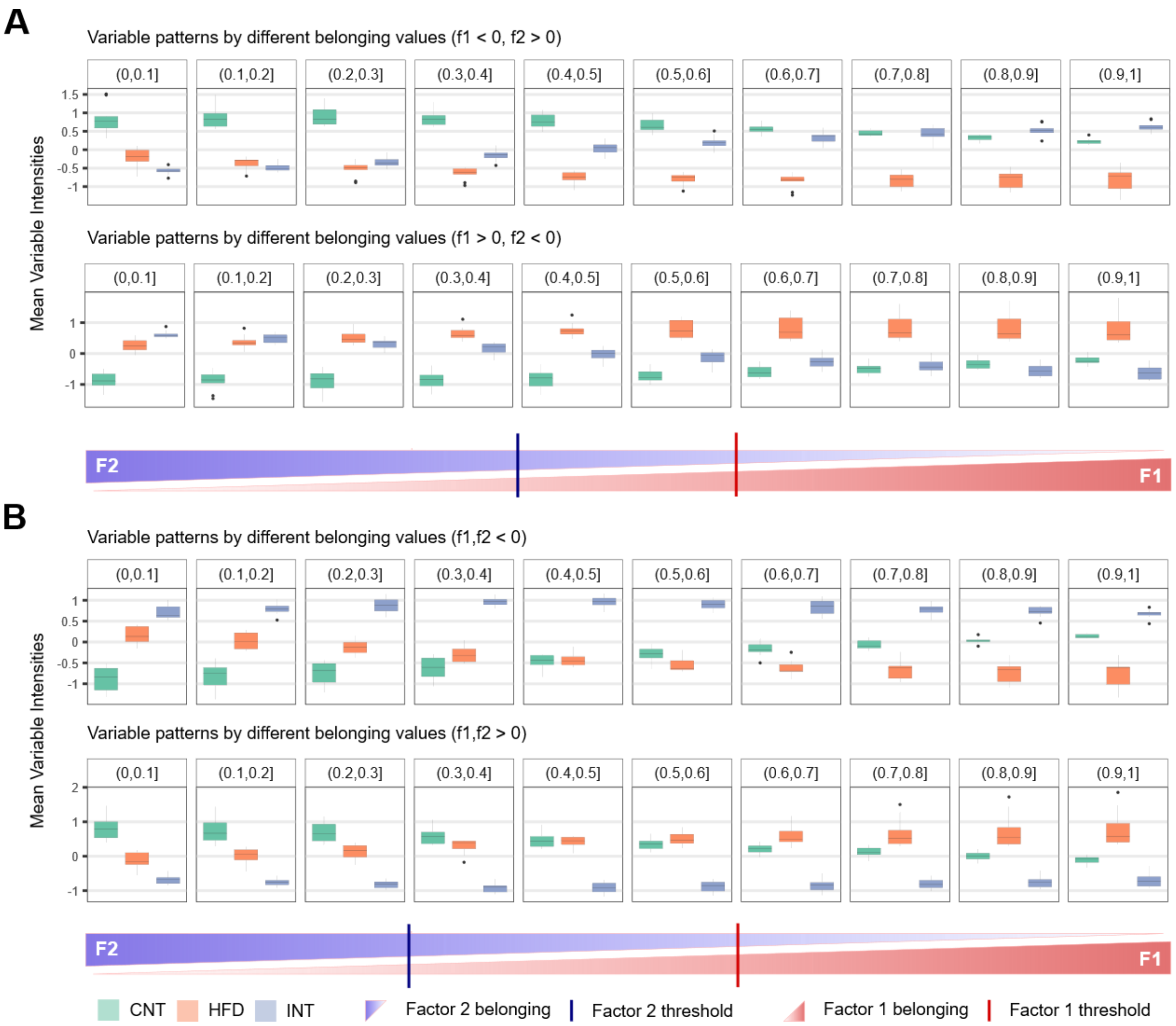

**Supplementary Figure 3**. **A.** Boxplots of the mean intensities of the variables with a given factor 1 belonging score interval (increasing from 0 to 1 in 0.1 intervals) for variables with opposite weights (f1 > 0 / f2 < 0 or f1 < 0 / f2 > 0) in the MOFA model. The lower bar represents the increasing belonging in factor 1 and decreasing in factor 2 and the blue and red marks indicate the thresholds selected for variable selection <0.4 for factor 2 and >0.6 for factor 1 respectively. **B.** Boxplots of the mean intensities of the variables with a given belonging score for variables with concordant weights (f1/f2 > 0 or f1/f2 < 0). The lower bar represents the increasing belonging in factor 1 and decreasing in factor 2 and the blue and red marks indicate the thresholds selected for variable selection <0.3 for factor 2 and >0.7 for factor 1 respectively.

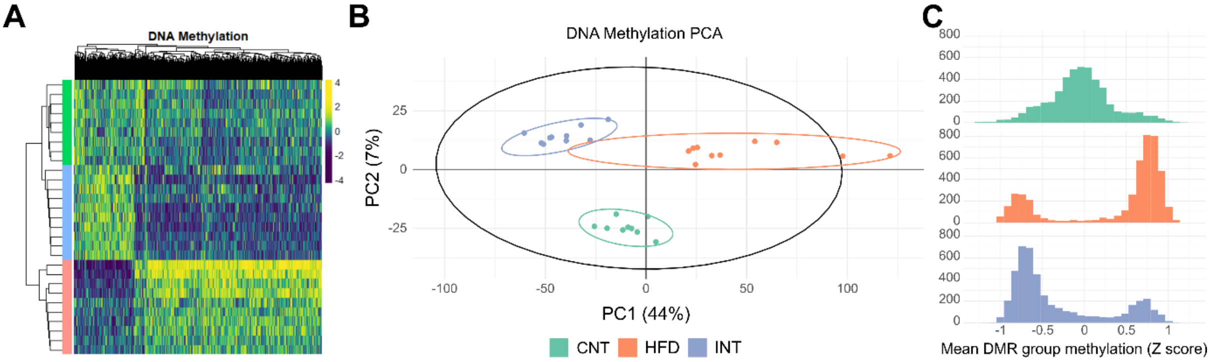


**Supplementary Figure 4. Genome-wide DNA methylation landscape across dietary conditions. A.** Heatmap of all differentially methylated regions (DMRs) identified by whole-genome bisulfite sequencing, with samples grouped by condition (CNT, control; HFD, high-fat diet; INT, lifestyle intervention) and DMRs ordered by hierarchical clustering. **B.** PCA of DMR methylation profiles. PC1 (44% of variance) separates HFD and INT animals at opposite poles, with control animals occupying an intermediate position, consistent with a directional methylation shift induced by HFD and further displaced by the intervention. PC2 (7% of variance) resolves a subset of DMRs that distinguishes control animals from both dietary groups, indicating that a fraction of HFD-induced methylation changes persist after intervention. **C.** Distribution of mean DMR methylation levels per group (VST Z-scores). Control animals display methylation centered around the population mean; HFD induces a rightward shift toward hypermethylation; the lifestyle intervention reverses this shift and displaces methylation below the control mean, suggesting active epigenetic remodeling beyond simple renormalization.

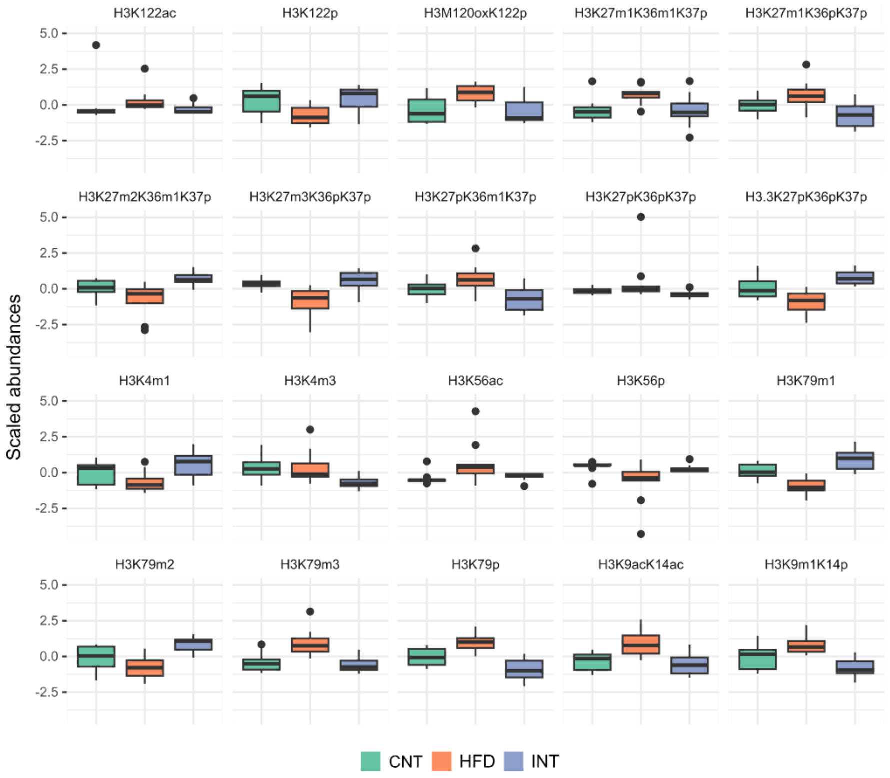


**Supplementary Figure 5**. **Histone H3 modifications across experimental groups.** Boxplots of the 20 histone H3 lysine modifications significantly altered by HFD, stratified by experimental group (control, HFD, and HFD followed by lifestyle intervention).

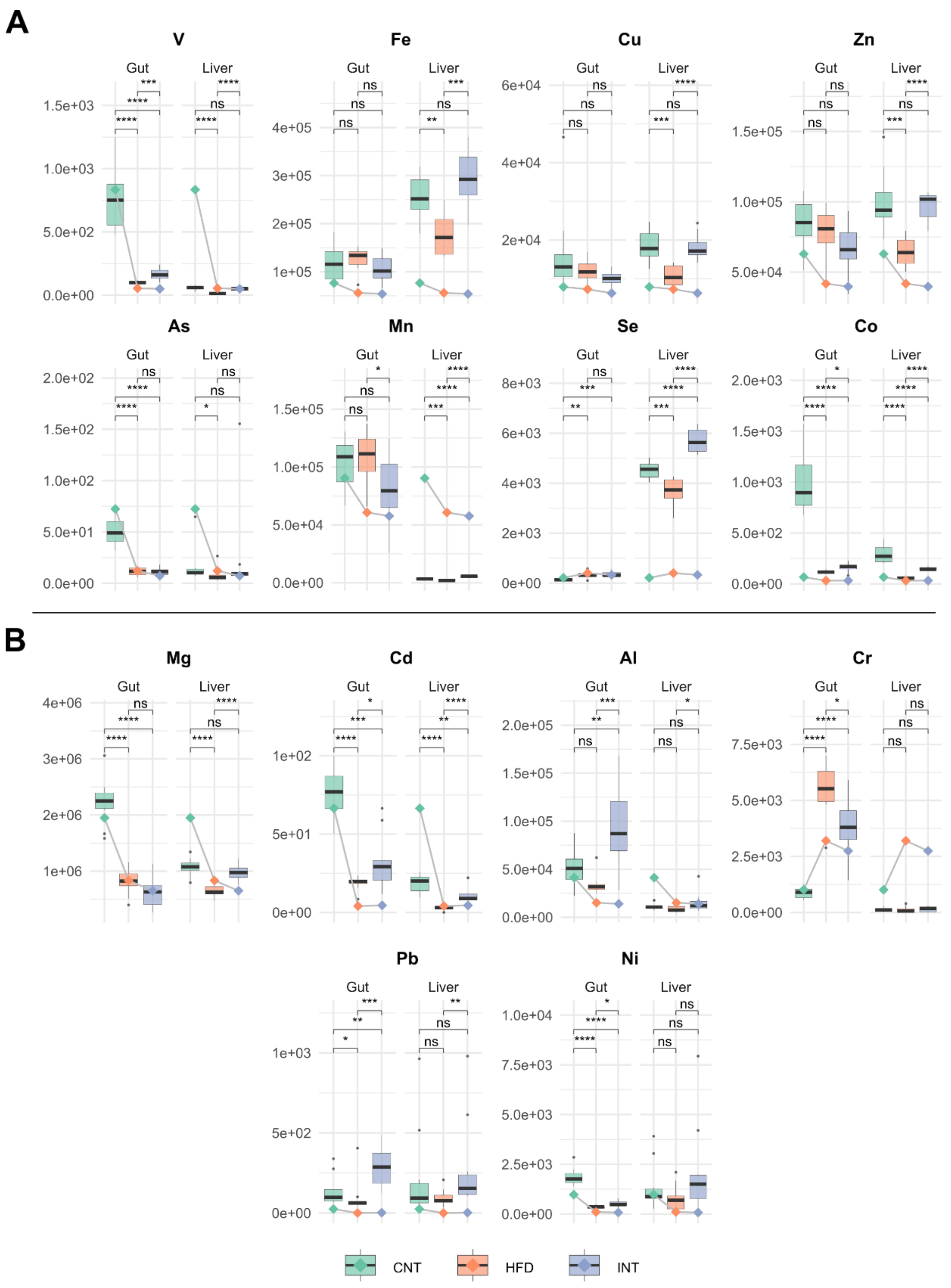


**Supplementary Figure 6**. **Metal abundances in gut and liver across experimental groups.** Boxplots show the abundance of 14 biologically relevant metals in gut and liver tissue, separated by experimental group (CNT, HFD, INT). Horizontal lines and dots indicate the corresponding metal content in the diet. **A.** Metals associated with the MOFA model: the seven metals contributing to Factor 1 (reversible response) and cobalt, the sole metal contributing to Factor 2 (persistent alterations). **B.** Metals that were not associated with any MOFA factor or showed no significant differences in the liver. (****p < 0.0001; ***p < 0.001; **p < 0.01; *p < 0.05; ns p > 0.05)

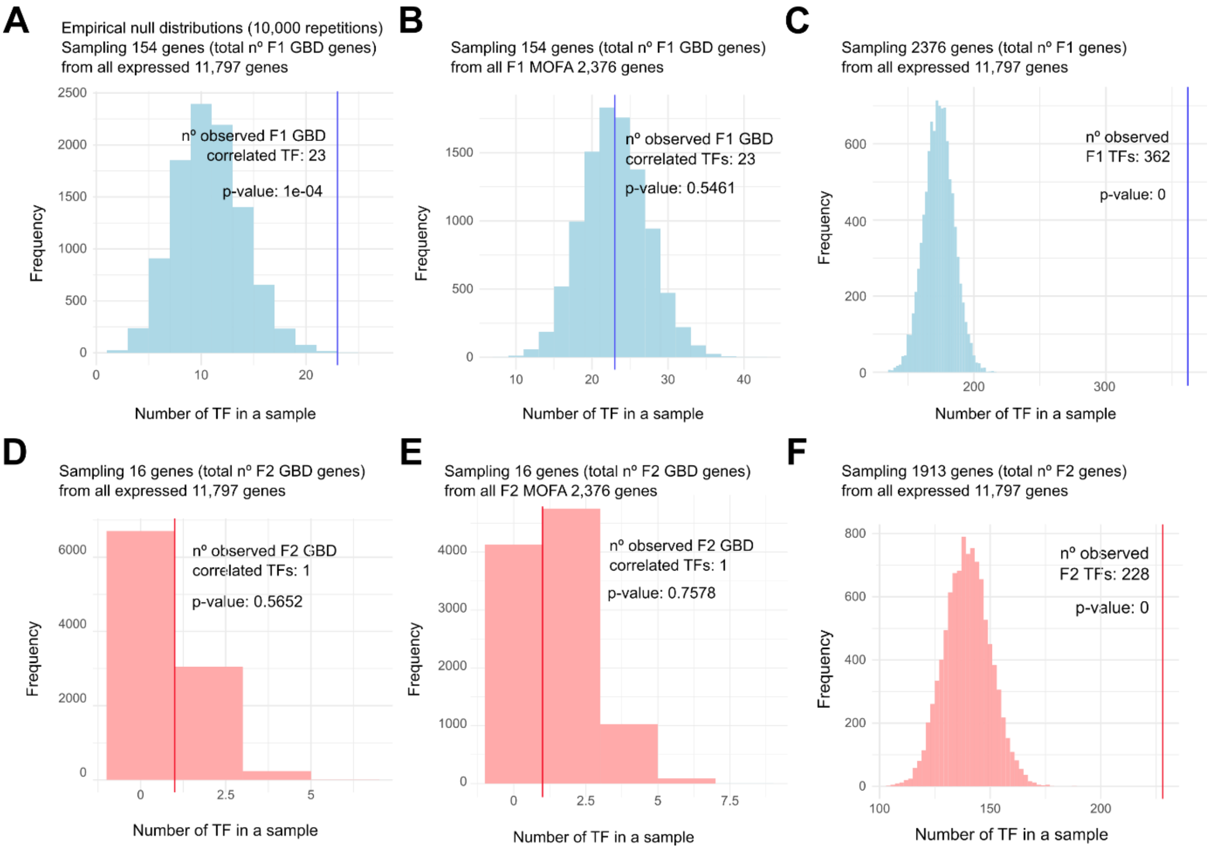


**Supplementary Figure 7. Empirical null distributions generated from permutation tests (10,000 random samples) assessing transcription factor (TF) enrichment**. **A.** Distribution of TF counts expected by chance from random sampling of all expressed genes (sample size = 154), compared with the observed number of factor 1 (F1) gene body DMR (GBD)-associated TFs (n = 23; ***p < 0.001). **B.** Distribution of TF counts expected by chance from random sampling of F1-associated genes (sample size = 154), compared with the observed number of F1 GBD-associated TFs (n = 23; ns, p > 0.05). **C.** Distribution of TF counts expected by chance from random sampling of all expressed genes (sample size = 2,376) compared with the observed number of F1-associated TFs (n = 362; ***p < 0.001). **D.** Distribution of TF counts expected by chance from random sampling of all expressed genes (sample size = 16), compared with the observed number of factor 2 (F2) GBD-associated TFs (n = 16; ns, p > 0.05). **E.** Distribution of TF counts expected by chance from random sampling of F2-associated genes (sample size = 16), compared with the observed number of F2 GBD-associated TFs (n = 1; ns, p > 0.05). **F.** Distribution of TF counts expected by chance from random sampling of all expressed genes (sample size = 1,913), compared with the observed total number of F2-associated TFs (n = 228; ***p < 0.001).

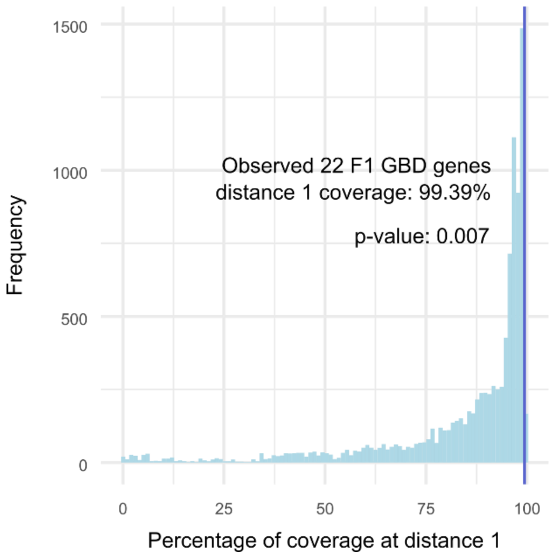


**Supplementary Figure 8. Empirical null distribution of TFLink network coverage of Factor 1 genes.** The empirical null distribution was generated by randomly sampling 22 TFs—the number of TFs associated with gene-body DMRs (GBD-regulated TFs) matched to TFLink—from the 348 TFs matched to the database, repeated 10,000 times. For each random set, the proportion of Factor 1 genes covered at network distance 1 was calculated. The observed coverage for the 22 GBD-regulated TFs is indicated by the vertical blue line (99.39%), which was significantly greater than expected by chance (permutation test, **p < 0.01).

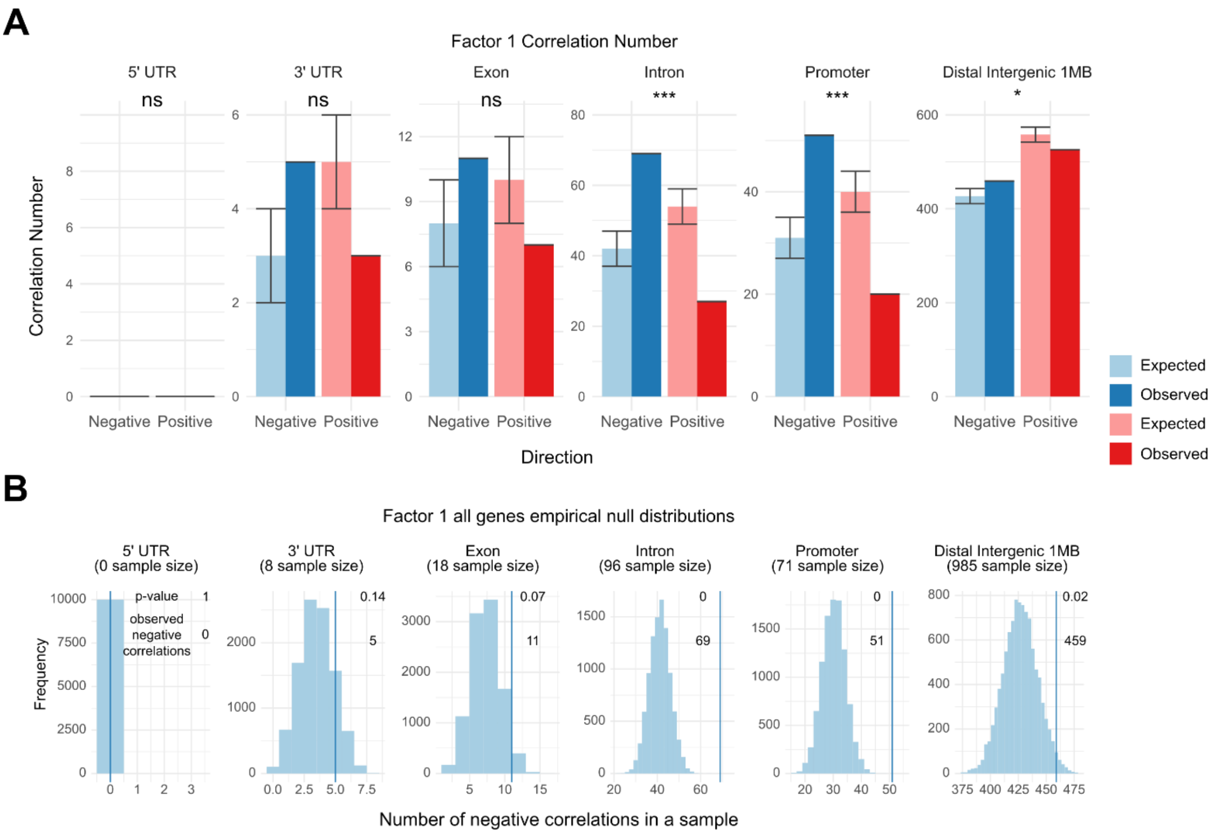


**Supplementary Figure 9. Directional bias of DMR–gene expression correlations in Factor 1. A.** Number of significant DMR–gene expression correlations (adj. p < 0.05) for Factor 1, grouped by DMR region, correlation direction (positive versus negative), and correlation type (observed versus expected). The expected number of correlations was derived by sampling from an empirical null distribution generated from all significant Factor 1 correlations. Statistical significance of the observed-versus-expected comparison: *p < 0.05; **p < 0.01; ***p < 0.001; ns, not significant. **B.** Empirical null distributions of the expected number of negative correlations, sampled from all significant Factor 1 correlations and shown separately by DMR region (corresponding to the comparisons in panel A). Dark blue vertical lines indicate the observed number of significant negative correlations for Factor 1.

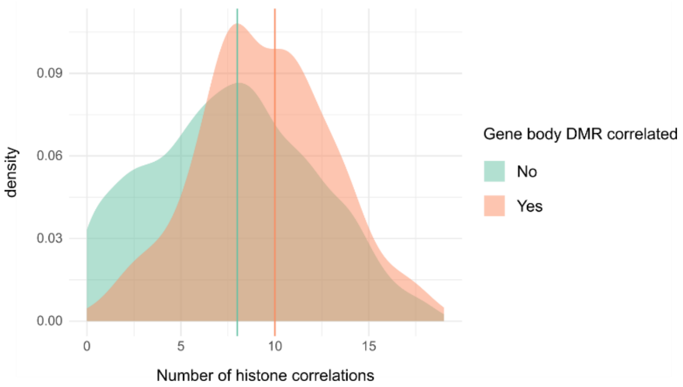


**Supplementary Figure 10. Genes with gene body DMRs show a higher number of histone modification correlations.** Density histogram showing the distribution of the number of significant gene–histone modification correlations (adj. p < 0.05) per gene, comparing genes harboring gene body DMRs with genes lacking them. The rightward shift of the gene body DMR-associated distribution indicates that these genes are correlated with a greater number of histone modifications than their non-DMR counterparts (****p < 0.0001).

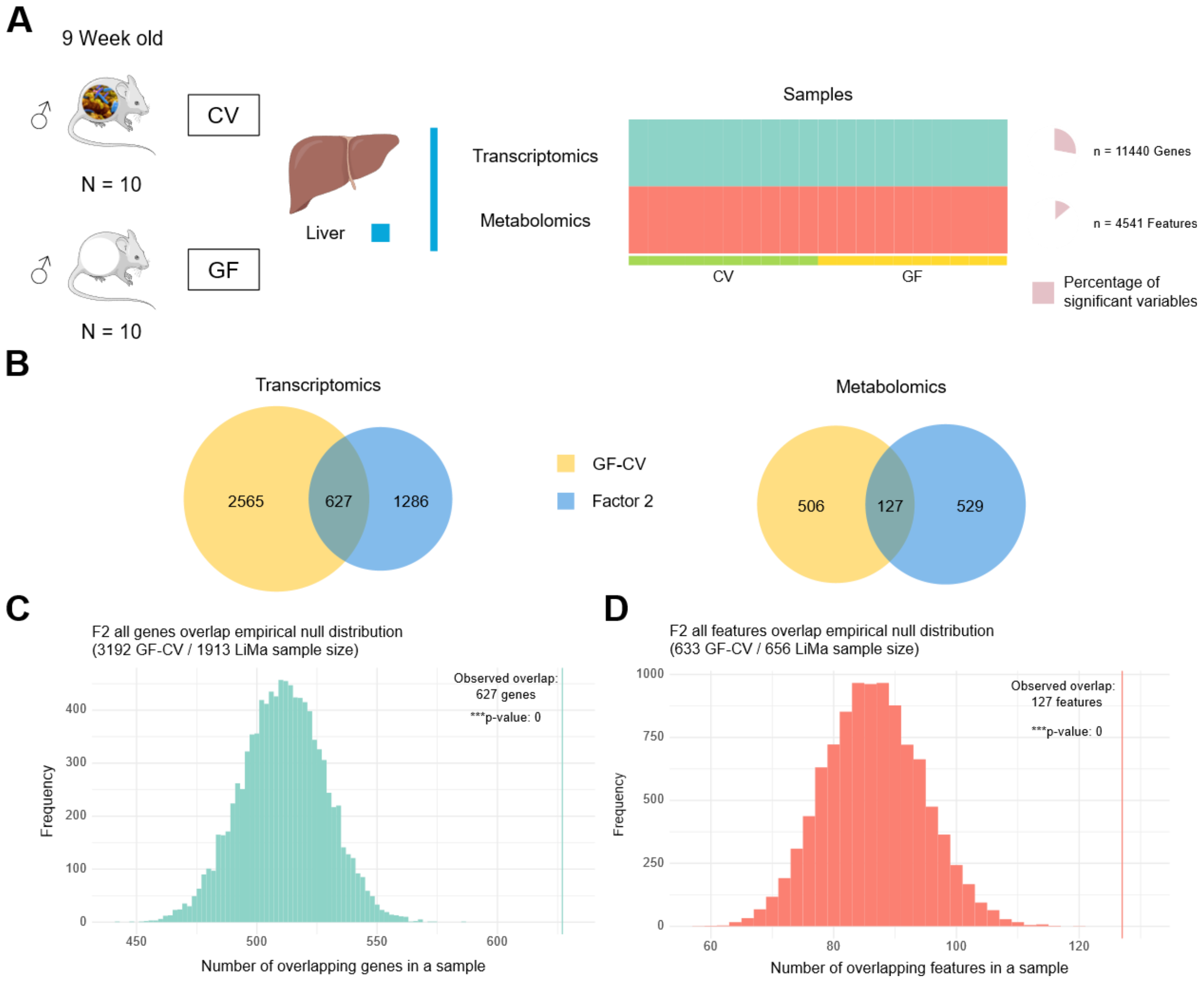


**Supplementary Figure 11. Definition of the microbiota-responsive hepatic space using germ-free mice. A.** Germ-free (GF) versus conventional (CV) experimental design. Nine-week-old black male germ-free mice and conventionally raised mice fed a standard chow diet (n = 10 per group) were compared to identify hepatic alterations dependent on microbial colonization. Liver transcriptomic and metabolomic profiles were acquired from both groups. The heatmap shows all measured variables across CV and GF samples; wedges on the right indicate the proportion of significantly different variables between groups (adj. $p < 0.05$), comprising 11,440 genes and 4,541 metabolite features in total. **B.** Venn diagrams showing the overlap between microbiota-responsive hepatic features (GF vs. CV differentially abundant variables, yellow) and the liver-associated features contributing to Factor 2 of the primary MOFA model (blue), shown separately for genes (left; 627 overlapping of 1,913 Factor 2 genes) and metabolite features (right; 127 overlapping of 656 Factor 2 features). The overlap defines the fraction of persistent Factor 2 liver alterations that fall within the microbiota-responsive hepatic space. **Permutation testing of the observed overlap between Factor 2 and microbiota-responsive hepatic features. C.** Null distribution for gene overlap. To assess whether the overlap between Factor 2 genes and GF–CV differentially expressed genes exceeded chance, genes were randomly sampled from the pool of shared LiMa and GF–CV genes using the observed set sizes (n = 1,913 Factor 2 genes; n = 3,192 GF–CV significant genes), and their overlap was computed. This procedure was repeated 10,000 times to generate the null distribution, which was compared to the observed overlap (n = 627; green line). The observed overlap significantly exceeded the null expectation (***$p < 0.001$, permutation test). **D.** Null distribution for metabolite overlap. The same permutation procedure was applied to metabolite features, using set sizes of n = 656 (Factor 2) and n = 633 (GF–CV), and the resulting null distribution was compared to the observed overlap (n = 127; red line). The observed overlap again significantly exceeded chance (***p < 0.001, permutation test).

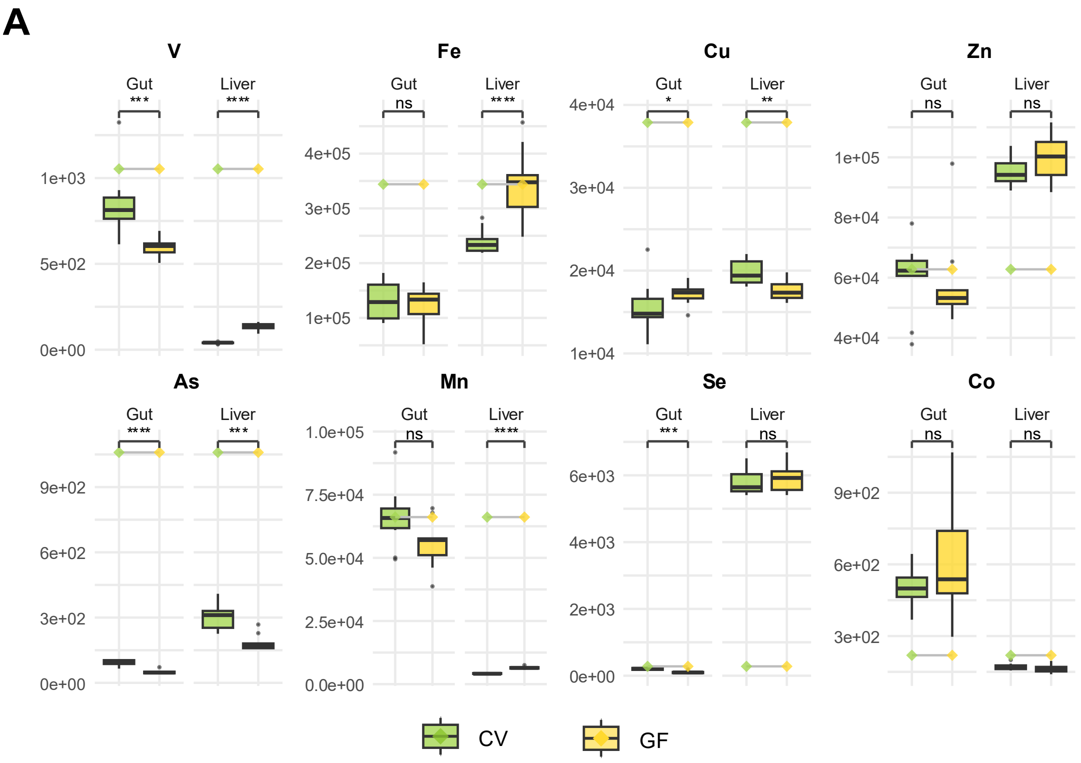


**Supplementary Figure 12**. Boxplots of biologically relevant metal abundances in the gut and liver from the Germ-Free model separated by experimental groups. Lines represent the quantity of the metal in the diet (****p < 0.0001; ***p < 0.001; **p < 0.01; *p < 0.05; ns p > 0.05).

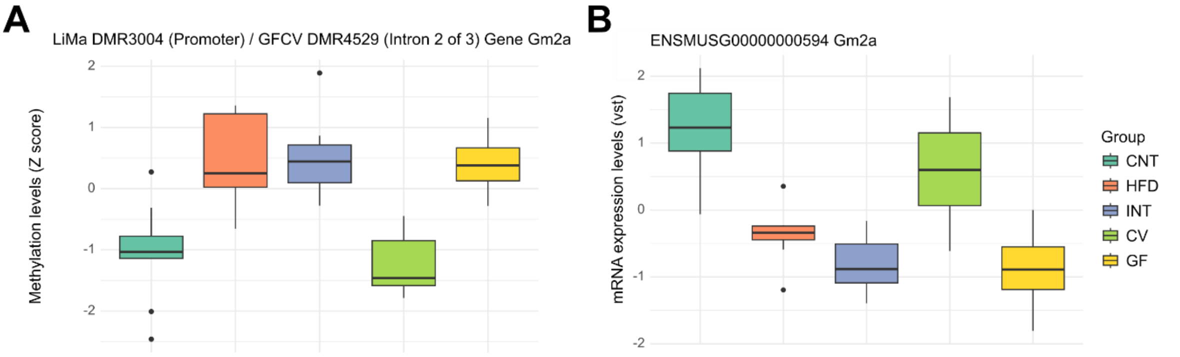


**Supplementary Figure 13. Gm2a DNA methylation and transcript abundance across experimental conditions. A.** Gm2a DNA methylation levels shown as Z scores. For LiMa samples (CNT, HFD and INT), methylation was measured at the Gm2a promoter DMR (DMR3004), whereas for GF-CV samples, methylation was measured in the second intron of Gm2a. **B.** Gm2a mRNA transcript abundance, shown as variance-stabilizing transformation (VST)-transformed counts, across the same experimental conditions.

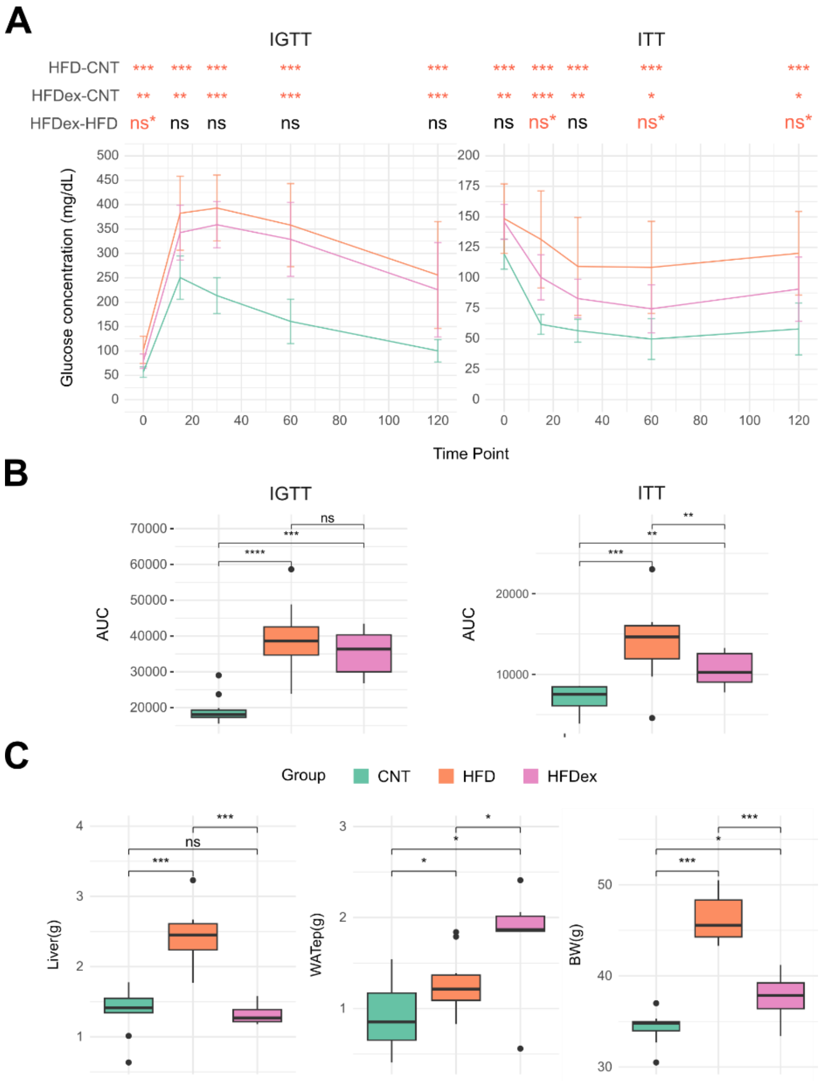


**Supplementary Figure 14. Metabolic characterization of low-responder (HFDex) mice. A.** Mean plasma glucose concentrations over time (0, 15, 30, 60, and 120 min) following glucose administration (intraperitoneal glucose tolerance test, IGTT; left) and insulin injection (insulin tolerance test, ITT; right), for control (CNT), high-fat diet (HFD), and HFD-excluded low-responder (HFDex) mice. Error bars represent standard deviation. Significance values for the pairwise contrasts HFD–CNT, HFDex–CNT, and HFDex–HFD are shown above each panel at the corresponding time points (FDR-adjusted p-values from a Conover post-hoc test following a significant Kruskal–Wallis test). **B.** Boxplots of the area under the curve (AUC) for the IGTT and ITT tests. **C.** Liver weight, epididymal white adipose tissue (WATep) weight, and body weight (BW) for CNT, HFD, and HFDex mice. Significance values correspond to FDR-corrected Wilcoxon test p-values. For all panels: ***p < 0.001; **p < 0.01; **p < 0.05; ns**, p < 0.1; ns, p > 0.1.

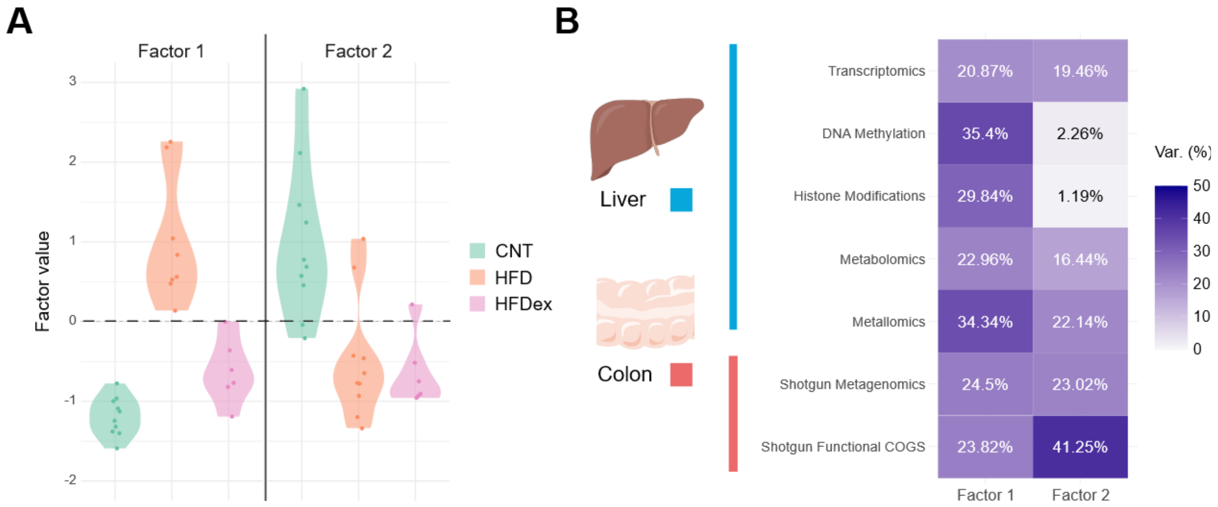


**Supplementary Figure 15. Second MOFA model incorporating low-responder (HFDex) mice. (A)** Violin plots of factor scores from the second MOFA model, in which the intervention group was replaced by low-responder (HFDex) mice, shown for Factor 1 and Factor 2 across the control (CNT), high-fat diet (HFD), and HFD-excluded (HFDex) groups. Factor 1 separates CNT (and, partially, HFDex) mice from HFD, capturing the low-responder phenotype, whereas Factor 2 separates both HFD and HFDex from CNT, capturing alterations shared with the full HFD phenotype. **(B)** Variance explained by each factor across the seven omic layers, measured in liver (transcriptomics, DNA methylation, histone modifications, metabolomics, metallomics) and colon (shotgun metagenomics and shotgun functional COG profiles). Consistent with the primary MOFA model, Factor 1 is predominantly driven by epigenetic (DNA methylation, histone modifications) and metallomic variables, while Factor 2 is primarily driven by colon microbial functional profiles (shotgun functional COGs, 41.25%).

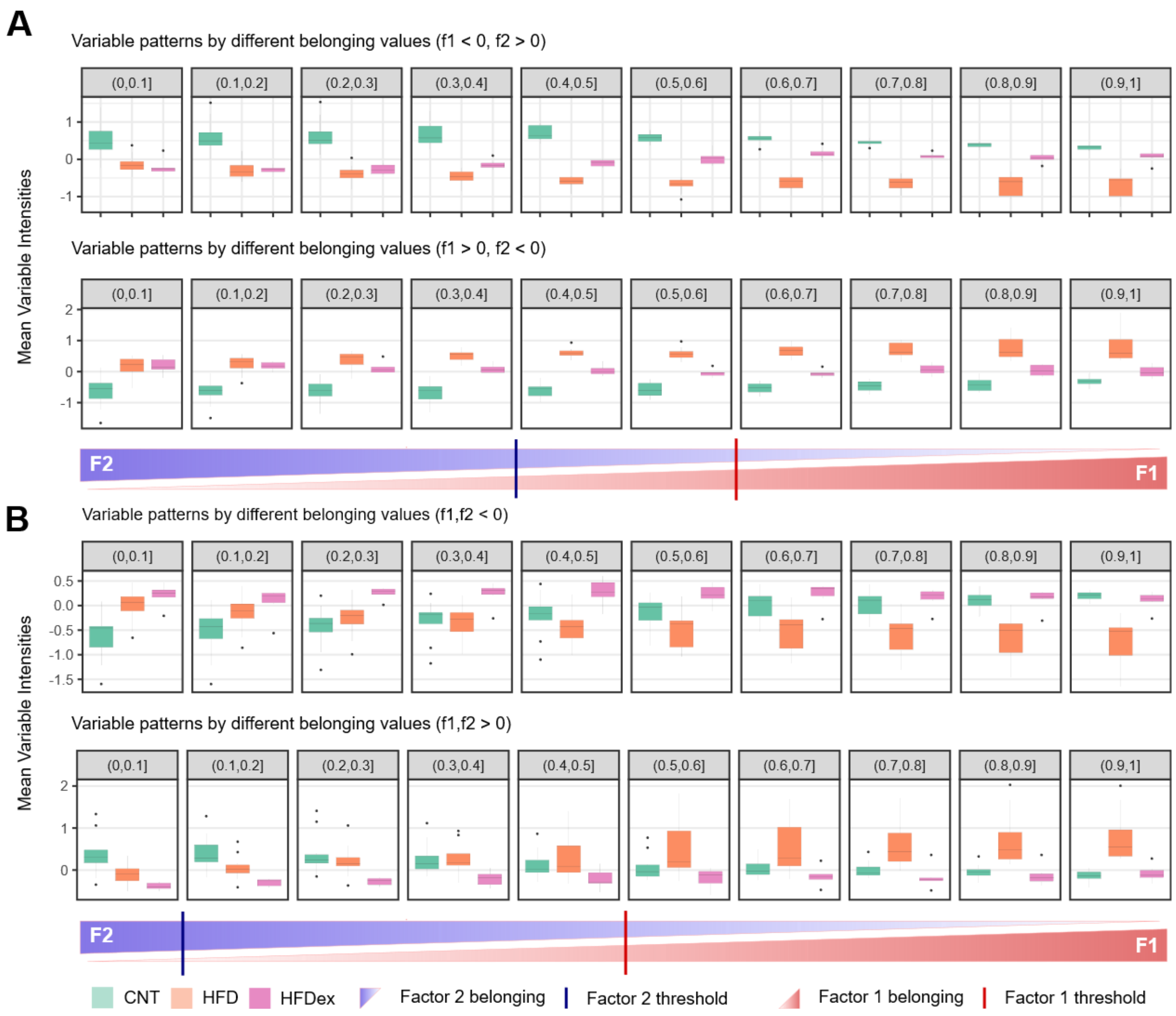


**Supplementary Figure 16. Weight-based variable selection for the second (HFDex) MOFA model.** For each variable, a belonging score reflects its relative assignment to Factor 1 versus Factor 2, based on the ratio of its MOFA weights; scores increase from 0 (fully assigned to Factor 2) to 1 (fully assigned to Factor 1). **(A)** Variables with opposite-sign weights across the two factors (F1 > 0 / F2 < 0, or F1 < 0 / F2 > 0). Boxplots show the mean intensities of variables within each belonging-score interval (0 to 1, in 0.1 increments); the horizontal axis reflects increasing Factor 1 assignment and decreasing Factor 2 assignment. Blue and red marks indicate the selection thresholds applied: < 0.4 for Factor 2 and > 0.6 for Factor 1, respectively. **(B)** Variables with concordant-sign weights across the two factors (F1 and F2 both > 0, or both < 0), plotted as in (A). Here the selection thresholds were < 0.1 for Factor 2 and > 0.5 for Factor 1.

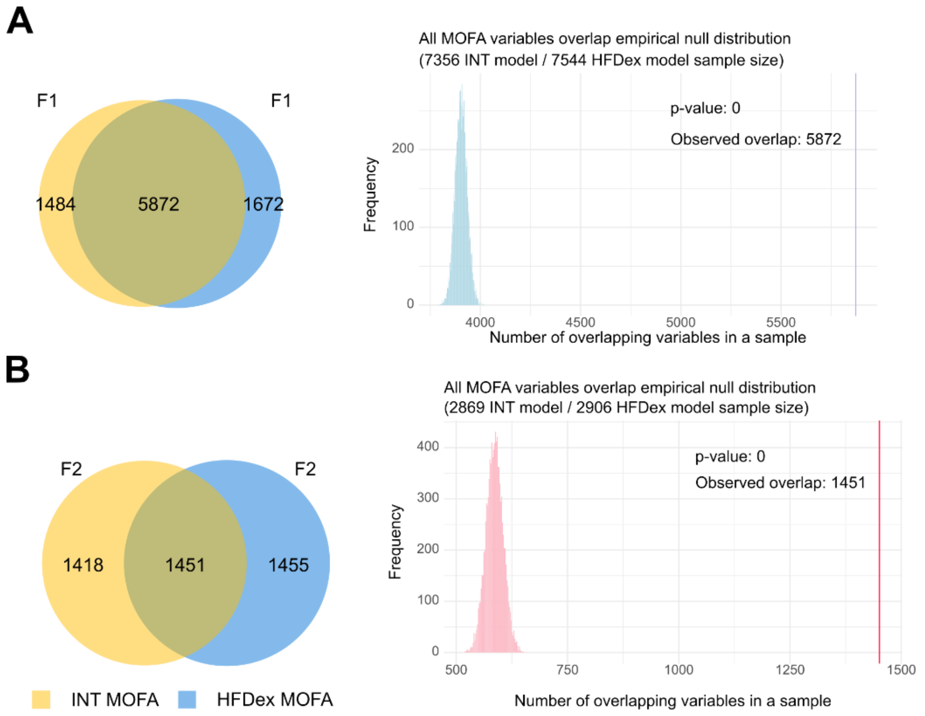


**Supplementary Figure 17. Overlap of factor-associated variables between the two MOFA models.** Venn diagrams (left) and permutation-based null distributions (right) comparing the variables assigned to each factor in the primary MOFA model (INT model, yellow) and the second MOFA model incorporating low-responder mice (HFDex model, blue). To assess whether the observed overlaps exceeded chance, X and Y variables (corresponding to the number of variables associated with the given factor in each model) were randomly sampled from the pool of variables common to both models, and their overlap computed. This was repeated 10,000 times to generate an empirical null distribution, which was compared to the observed overlap. **(A)** Factor 1 variables from both models. The Venn diagram shows 5,872 shared variables (1,484 unique to the INT model, 1,672 unique to the HFDex model), corresponding to ~80% overlap. The observed overlap (blue line; n = 5,872) far exceeded the null distribution (sample sizes: 7,356 INT, 7,544 HFDex; ****p < 0.0001, permutation test). **(B)** Factor 2 variables from both models. The Venn diagram shows 1,451 shared variables (1,418 unique to INT, 1,455 unique to HFDex), corresponding to ~50% overlap. The observed overlap (red line; n = 1,451) again far exceeded the null distribution (sample sizes: 2,869 INT, 2,906 HFDex; ****p < 0.0001, permutation test)

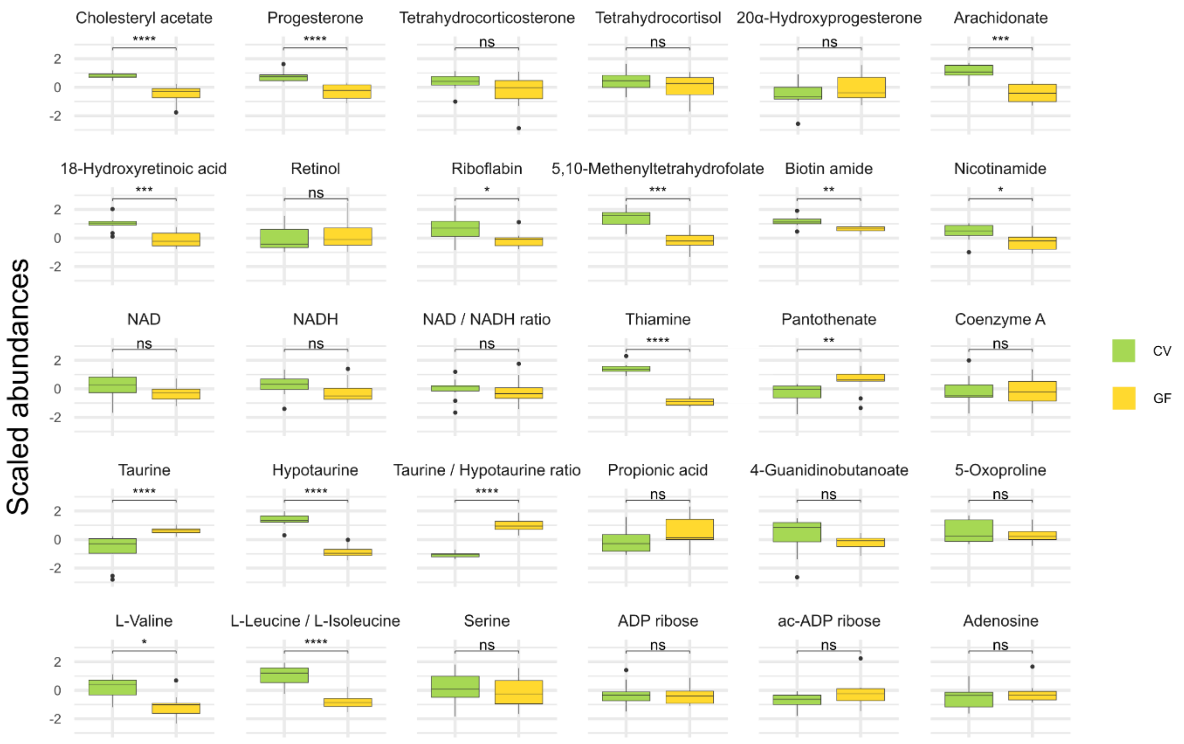


**Supplementary Figure 18. Microbiota dependence of early hepatic metabolite alterations, assessed in germ-free versus conventional mice.** Hepatic abundances of the same representative metabolites shown in Figure 6B, spanning steroid and cholesterol metabolism, retinol metabolism, vitamin and cofactor metabolism (thiamine, riboflavin, biotin, pantothenate, nicotinamide, folate derivatives, NAD/NADH, CoA), sulfur amino acids (taurine and hypotaurine metabolism), propanoate metabolism, and branched-chain amino acid degradation, compared between conventionally raised (CV, green) and germ-free (GF, yellow) mice. Boxplots show scaled metabolite abundances; significance: *p < 0.05; **p < 0.01; ***p < 0.001; ****p < 0.0001; ns p > 0.05

**Supplementary Table 1.** Diet metal concentrations (ng/g)

| (ng/g) | Mg | Al | V | Cr | Mn | Fe | Co | Ni | Cu | Zn | As | Se | Cd | Pb |
|---|---|---|---|---|---|---|---|---|---|---|---|---|---|---|
| CTR | 1,949,525 | 41,229 | 834 | 1013 | 90,323 | 76,608 | 66 | 973 | 7883 | 62,931 | 72 | 211 | 66 | 25 |
| HFD | 833,269 | 15,149 | 55 | 3198 | 60,699 | 56,153 | 31 | 103 | 7278 | 41,748 | 12 | 399 | 4.1 | <0.01 |
| INT | 650,664 | 13,977 | 50 | 2747 | 57,620 | 53,947 | 32 | 75 | 6263 | 39,718 | 7.3 | 328 | 4.5 | 2.2 |
| GF/CV | 2,226,208 | 144,299 | 1053 | 852 | 66,132 | 344,074 | 220 | 10,560 | 37,889 | 62,776 | 1059 | 280 | 76 | 119 |

**Supplementary Table 2.** Gene body DMRs encoding transcription factors from factor 1.

| Ensembl ID | Entrez ID | Symbol | Gene Name | Chromosome |
|---|---|---|---|---|
| ENSMUSG00000002881 | 17936 | Nab1 | Ngfi-A binding protein 1 | chr1 |
| ENSMUSG00000026398 | 26424 | Nr5a2 | nuclear receptor subfamily 5, group A, member 2 | chr1 |
| ENSMUSG00000002111 | 20375 | Spi1 | spleen focus forming virus (SFFV) proviral integration oncogene | chr2 |
| ENSMUSG00000032698 | 16909 | Lmo2 | LIM domain only 2 | chr2 |
| ENSMUSG00000027177 | 15259 | Hipk3 | homeodomain interacting protein kinase 3 | chr2 |

| | | | | |
|---|---|---|---|---|
| ENSMUSG00000017950 | 15378 | Hnf4a | hepatic nuclear factor 4, Alpha | chr2 |
| ENSMUSG00000039852 | 68703 | Rere | arginine glutamic acid dipeptide (RE) repeats | chr4 |
| ENSMUSG00000029635 | 264064 | Cdk8 | cyclin dependent kinase 8 | chr5 |
| ENSMUSG00000009376 | 17295 | Met | met proto-oncogene | chr6 |
| ENSMUSG00000051910 | 20679 | Sox6 | SRY (sex determining region Y)-box 6 | chr7 |
| ENSMUSG00000054717 | 97165 | Hmgb2 | high mobility group box 2 | chr8 |
| ENSMUSG00000041438 | 21771 | Utp4 | UTP4 small subunit processome component | chr8 |
| ENSMUSG00000019947 | 71371 | Arid5b | AT-rich interaction domain 5B | chr10 |
| ENSMUSG00000018654 | 22778 | Ikzf1 | IKAROS family zinc finger 1 | chr11 |
| ENSMUSG00000037149 | 104721 | Ddx1 | DEAD box helicase 1 | chr12 |
| ENSMUSG00000021258 | 12454 | Ccnk | cyclin K | chr12 |
| ENSMUSG00000021457 | 20963 | Syk | spleen tyrosine kinase | chr13 |
| ENSMUSG00000006527 | 54650 | Sfmbt1 | Scm-like with four mbt domains 1 | chr14 |
| ENSMUSG00000033565 | 93686 | Rbfox2 | RNA binding protein, fox-1 homolog (C. elegans) 2 | chr15 |
| ENSMUSG00000023852 | 12648 | Chd1 | chromodomain helicase DNA binding protein 1 | chr17 |
| ENSMUSG00000002249 | 21678 | Tead3 | TEA domain family member 3 | chr17 |
| ENSMUSG00000023951 | 22339 | Vegfa | vascular endothelial growth factor A | chr17 |
| ENSMUSG00000042439 | 328977 | Zfp532 | zinc finger protein 532 | chr18 |

**Supplementary Table 3. Taxonomy and control-group abundance of the 14 reversible genera associated with Factor 1.** For each of the 14 genera contributing to Factor 1, the table lists the full taxonomic classification, the mean absolute abundance in control (CNT) samples, and the corresponding relative abundance expressed as a percentage of total genus-level counts (unfiltered). Older taxonomy identifications were updated, and changes are available at the zenodo.

| Phylum | Class | Order | Family | Genus | Mean CNT Abundance | Mean CNT abundance percentage |
|---|---|---|---|---|---|---|
| Bacteroidota | Bacteroidia | Bacteroidales | Bacteroidaceae | Bacteroides | 33 | 1.73 |
| Bacteroidota | Bacteroidia | Bacteroidales | Odoribacteraceae | Odoribacter | 44.2 | 2.32 |
| Bacteroidota | Bacteroidia | Bacteroidales | Prevotellaceae | Alloprevotella | 13.6 | 0.71 |
| Bacillota | Clostridia | Eubacteriales | Oscillospiraceae | Intestinimonas | 15.2 | 0.80 |
| Bacillota | Clostridia | Eubacteriales | Clostridiaceae | Clostridium | 3.4 | 0.18 |
| Bacillota | Clostridia | Eubacteriales | Erysipelotrichaceae | Ileibacterium | 13.6 | 0.71 |
| Bacillota | Clostridia | Eubacteriales | Lachnospiraceae | Lachnoclostridium | 32.2 | 1.69 |
| Bacillota | Clostridia | Eubacteriales | Lachnospiraceae | Roseburia | 60.2 | 3.16 |
| Bacillota | Clostridia | Eubacteriales | Oscillospiraceae | Oscillibacter | 24.5 | 1.67 |
| Bacillota | Erysipelotrichia | Erysipelotrichales | Erysipelotrichaceae | Faecalitalea | 1.4 | 0.07 |
| Deferribacterota | Deferribacteres | Deferribacterales | Deferribacteraceae | Mucispirillum | 14.9 | 1.08 |
| Bacillota | Bacilli | Lactobacillales | Lactobacillaceae | Lactobacillus | 310.8 | 16.35 |
| Bacillota | Clostridia | Eubacteriales | Ruminococcaceae | Ruminiclostridium | 30.2 | 1.59 |
| Pseudomonadota | Deltaproteobacteria | Desulfovibrionales | Desulfovibrionaceae | Bilophila | 5.8 | 0.30 |

**Supplementary Table 4. COG functionalities and associated functional pathways of the 14 Factor 1 reversible genera.** The table lists the 48 Clusters of Orthologous Groups (COG) functional terms encoded by the 14 genera associated with Factor 1, together with the 17 higher-order functional pathways to which they map.

| COG ID | Gene | Description | Associated Functional Pathway |
|---|---|---|---|
| COG0049 | RpsG | Ribosomal protein S7 | Ribosome 30S subunit |
| COG0157 | NadC | Nicotinate-nucleotide pyrophosphorylase | NAD biosynthesis |
| COG0178 | UvrA | Excinuclease UvrABC ATPase subunit | |
| COG0312 | TldD | Zn-dependent protease or N-deacetylase, PmbA/TldD/TldE family | |
| COG0522 | RpsD | Ribosomal protein S4 or related protein | Ribosome 30S subunit |
| COG0776 | HupA | DNA-binding chromatin protein HU or IHF, alpha or beta variants | |
| COG1592 | YotD | Rubrerythrin | |
| COG2156 | KdpC | K+-transporting ATPase, KdpC subunit | |
| COG2826 | Tra8 | Transposase and inactivated derivatives, IS30 family | |
| COG3039 | IS5 | Transposase and inactivated derivatives, IS5 family | |
| COG3385 | InsG | IS4 transposase InsG | |
| COG4799 | MmdA | Acetyl-CoA carboxylase, carboxyltransferase component | |
| COG4974 | XerD | Site-specific tyrosine recombinase XerD | |
| COG0089 | RplW | Ribosomal protein L23 | Ribosome 50S subunit |
| COG0504 | PyrG | CTP synthase (UTP-ammonia lyase) | Pyrimidine biosynthesis |
| COG0771 | MurD | UDP-N-acetylmuramoylalanine-D-glutamate ligase | Mureine biosynthesis |
| COG1733 | HxlR | DNA-binding transcriptional regulator, HxlR family | |
| COG3250 | LacZ | Beta-galactosidase/beta-glucuronidase | |
| COG0169 | AroE | Shikimate 5-dehydrogenase | Aromatic amino acid biosynthesis |
| COG0228 | RpsP | Ribosomal protein S16 | Ribosome 30S subunit |
| COG0352 | ThiE | Thiamine monophosphate synthase | Thiamine biosynthesis |
| COG0413 | PanB | Ketopantoate hydroxymethyltransferase | Pantothenate/CoA biosynthesis |
| COG0634 | HptA | Hypoxanthine-guanine phosphoribosyltransferase | Purine salvage |
| COG0720 | QueD | 6-pyruvoyl-tetrahydropterin synthase | tRNA modification |
| COG0781 | NusB | Transcription antitermination protein NusB | |
| COG1092 | RlmK | 23S rRNA G2069 N7-methylase RlmK or C1962 C5-methylase RlmI | 23S rRNA modification |
| COG1115 | AlsT | Na+/alanine symporter | |
| COG1158 | Rho | Transcription termination factor Rho | |
| COG1191 | FliA | DNA-directed RNA polymerase specialized sigma subunit | RNA polymerase |
| COG1347 | NqrD | Na+-transporting NADH:ubiquinone oxidoreductase, subunit NqrD | Na+-translocating NADH dehydrogenase |
| COG1670 | RimL | Protein N-acetyltransferase, RimJ/RimL family | |
| COG1884 | Sbm1 | Methylmalonyl-CoA mutase, N-terminal domain/subunit | |
| COG3341 | Rnh1 | Ribonuclease HI-related protein, contains viroplasmin and RNaseH domains | |
| COG3344 | YkfC | Retron-type reverse transcriptase | |
| COG3426 | Buk | Butyrate kinase | |
| COG3682 | CopY | Transcriptional regulator, CopY/TcrY family | |
| COG4115 | | Toxin component of the Txe-Axe toxin-antitoxin module, Txe/YoeB family | |

| | | | |
|---|---|---|---|
| COG4123 | TrmN6 | tRNA1(Val) A37 N6-methylase TrmN6 | tRNA modification |
| COG4658 | RnfD | Na+-translocating ferredoxin:NAD+ oxidoreductase RNF, RnfD subunit | Na+-translocating Fd:NADH oxidoreductase |
| COG4804 | YhcG | Predicted nuclease of restriction endonuclease-like (RecB) superfamily, DUF1016 family | |
| COG0154 | GatA | Asp-tRNAAsn/Glu-tRNAGln amidotransferase A subunit or related amidase | Aminoacyl-tRNA synthetases |
| COG0223 | Fmt | Methionyl-tRNA formyltransferase | |
| COG0345 | ProC | Pyrroline-5-carboxylate reductase | Proline biosynthesis |
| COG0671 | PgpB | Membrane-associated phospholipid phosphatase | Phospholipid biosynthesis |
| COG1176 | PotB | ABC-type spermidine/putrescine transport system, permease component I | |
| COG1354 | ScpA | Chromatin segregation and condensation protein Rec8/ScpA/Scc1, kleisin family | |
| COG3331 | YotM | Penicillin-binding protein-related factor A, putative recombinase | |
| COG4988 | CydD | ABC-type transport system involved in cytochrome bd biosynthesis, ATPase and permease components | |

**Supplementary Table 5. Taxonomy and control-group abundance of the seven persistent genera associated with Factor 2.** For each of the 7 genera contributing to Factor 2, the table lists the full taxonomic classification, the mean absolute abundance in control (CNT) samples, and the corresponding relative abundance expressed as a percentage of total genus-level counts (unfiltered). Older taxonomy identifications were updated, and changes are available at the zenodo.

| Phylum | Class | Order | Family | Genus | Mean CNT abundance | Mean CNT abundance (%) |
|---|---|---|---|---|---|---|
| Bacillota | Bacilli | Lactobacillales | Streptococcaceae | Lactococcus | 0 | 0 |
| Bacillota | Clostridia | Eubacteriales | Lachnospiraceae | Acetatifactor | 16.6 | 0.874 |
| Bacillota | Clostridia | Eubacteriales | Lachnospiraceae | Eisenbergiella | 5.2 | 0.274 |
| Bacillota | Clostridia | Eubacteriales | Lachnospiraceae | Marvinbryantia | 10 | 0.526 |
| Bacillota | Clostridia | Eubacteriales | Oscillospiraceae | Ruminococcus | 11.6 | 0.610 |
| Proteobacteria | Betaproteobacteria | Burkholderiales | Sutterellaceae | Parasutterella | 7.8 | 0.410 |
| Tenericutes | Mollicutes | Anaeroplasmatales | Anaeroplasmataceae | Anaeroplasma | 22.4 | 1.179 |

**Supplementary Table 6. COG functionalities and associated functional pathways of the seven Factor 2 persistent genera.** The table lists the 170 Clusters of Orthologous Groups (COG) functional terms encoded by the 7 genera associated with Factor 2, together with the 34 higher-order functional pathways to which they map.

| COG ID | Gene | Description | Associated Functional Pathway |
|---|---|---|---|
| COG0087 | RplC | Ribosomal protein L3 | Ribosome 50S subunit |
| COG0092 | RpsC | Ribosomal protein S3 | Ribosome 30S subunit |
| COG0094 | RplE | Ribosomal protein L5 | Ribosome 50S subunit |
| COG0142 | IspA | Geranylgeranyl pyrophosphate synthase | Isoprenoid biosynthesis |
| COG0148 | Eno | Enolase | Glycolysis |
| COG0201 | SecY | Preprotein translocase subunit SecY | Sec pathway |
| COG0359 | RplI | Ribosomal protein L9 | Ribosome 50S subunit |
| COG0669 | CoaD | Phosphopantetheine adenylyltransferase | Pantothenate/CoA biosynthesis |
| COG0840 | Tar | Methyl-accepting chemotaxis protein (MCP) | |
| COG1294 | AppB | Cytochrome bd-type quinol oxidase, subunit 2 | |

| | | | |
|---|---|---|---|
| COG1501 | YicI | Alpha-glucosidase/xylosidase, GH31 family | |
| COG1595 | RpoE | DNA-directed RNA polymerase specialized sigma subunit, sigma24 family | RNA polymerase |
| COG1739 | YIH1 | Putative translation regulator, IMPACT (imprinted ancient) protein family | |
| COG2115 | XylA | Xylose isomerase | |
| COG3104 | PTR2 | Dipeptide/tripeptide permease | |
| COG3507 | XynB2 | Beta-xylosidase | |
| COG4537 | ComGC | Competence protein ComGC | |
| COG0018 | ArgS | Arginyl-tRNA synthetase | Aminoacyl-tRNA synthetases |
| COG0098 | RpsE | Ribosomal protein S5 | Ribosome 30S subunit |
| COG0193 | Pth | Peptidyl-tRNA hydrolase | Translation factors |
| COG0197 | RplP | Ribosomal protein L16/L10AE | Ribosome 50S subunit |
| COG0511 | AccB | Biotin carboxyl carrier protein | Fatty acid biosynthesis |
| COG0513 | SrmB | Superfamily II DNA and RNA helicase | |
| COG0618 | NrnA | nanoRNase/pAp phosphatase, hydrolyzes c-di-AMP and oligoRNAs | |
| COG0804 | UreC | Urease alpha subunit | |
| COG0820 | RlmN | Adenine C2-methylase RlmN of 23S rRNA A2503 and tRNA A37 | 23S rRNA modification |
| COG1190 | LysU | Lysyl-tRNA synthetase, class II | Aminoacyl-tRNA synthetases |
| COG1234 | ElaC | Ribonuclease BN, tRNA processing enzyme | |
| COG1312 | UxuA | D-mannonate dehydratase | |
| COG1674 | FtsK | DNA segregation ATPase FtsK/SpoIIIE or related protein | |
| COG2942 | YihS | Mannose or cellobiose epimerase, N-acyl-D-glucosamine 2-epimerase family | |
| COG3534 | AbfA | Alpha-L-arabinofuranosidase | |
| COG3855 | FbpC | Fructose-1,6-bisphosphatase | Gluconeogenesis |
| COG0006 | PepP | Xaa-Pro aminopeptidase | |
| COG0029 | NadB | Aspartate oxidase | NAD biosynthesis |
| COG0053 | FieF | Divalent metal cation (Fe/Co/Zn/Cd) efflux pump | |
| COG0073 | EMAP | tRNA-binding EMAP/Myf domain | |
| COG0077 | PheA2 | Prephenate dehydratase (decarboxylase) | Aromatic amino acid biosynthesis |
| COG0132 | BioD | Dethiobiotin synthetase | Biotin biosynthesis |
| COG0156 | BioF | 7-keto-8-aminopelargonate synthetase or related enzyme | Biotin biosynthesis |
| COG0161 | BioA | Adenosylmethionine-8-amino-7-oxononanoate aminotransferase | Biotin biosynthesis |
| COG0195 | NusA | Transcription antitermination factor NusA, contains S1 and KH domains | |
| COG0232 | Dgt | dGTP triphosphohydrolase | |
| COG0234 | GroES | Co-chaperonin GroES (HSP10) | |
| COG0246 | MtlD | Mannitol-1-phosphate/altronate dehydrogenases | |
| COG0247 | GlpC | Fe-S cluster-containing oxidoreductase, includes glycolate oxidase subunit GlcF | |
| COG0270 | Dcm | DNA-cytosine methylase Dcm or eukaryotic tRNA-C38 C5-methylase, Dcm/DNMT2/TRDMT1 family | |
| COG0285 | FolC | Folylpolyglutamate synthase/Dihydropteroate synthase | Folate biosynthesis |
| COG0302 | FolE | GTP cyclohydrolase I | Folate biosynthesis |
| COG0339 | Dcp | Zn-dependent oligopeptidase, M3 family | |
| COG0340 | BirA2 | Biotin-protein ligase | Biotin biosynthesis |

| | | | |
|---|---|---|---|
| COG0341 | SecF | Preprotein translocase subunit SecF | |
| COG0365 | Acs | Acyl-coenzyme A synthetase/AMP-(fatty) acid ligase | |
| COG0368 | CobS | Cobalamin synthase CobS (adenosylcobinamide-GDP ribazoletransferase) | Cobalamine/B12 biosynthesis |
| COG0377 | NuoB | NADH:ubiquinone oxidoreductase 20 kD subunit (chain B) or related Fe-S oxidoreductase | NADH dehydrogenase |
| COG0379 | NadA | Quinolinate synthase | NAD biosynthesis |
| COG0414 | PanC | Panthothenate synthetase | Pantothenate/CoA biosynthesis |
| COG0422 | ThiC | 4-amino-2-methyl-5-hydroxymethylpyrimidine (HMP) synthase ThiC | Thiamine biosynthesis |
| COG0427 | ACH1 | Propionyl CoA:succinate CoA transferase | |
| COG0445 | MnmG | tRNA U34 5-carboxymethylaminomethyl modifying enzyme MnmG/GidA | tRNA modification |
| COG0450 | AhpC | Alkyl hydroperoxide reductase subunit AhpC (peroxiredoxin) | |
| COG0476 | ThiF | Molybdopterin or thiamine biosynthesis adenylyltransferase | Molybdopterin biosynthesis |
| COG0483 | SuhB | Archaeal fructose-1,6-bisphosphatase or related enzyme, inositol monophosphatase family | Gluconeogenesis |
| COG0496 | SurE | Broad specificity polyphosphatase and 5'/3'-nucleotidase SurE | |
| COG0499 | SAM1 | S-adenosylhomocysteine hydrolase | |
| COG0502 | BioB | Biotin synthase or related enzyme | Biotin biosynthesis |
| COG0517 | CBS | CBS domain | |
| COG0523 | YejR | Zinc metallochaperone YeiR/ZagA and related GTPases, G3E family | |
| COG0525 | ValS | Valyl-tRNA synthetase | Aminoacyl-tRNA synthetases |
| COG0535 | SkfB | Radical SAM superfamily maturase, SkfB/NifB/PqqE family | |
| COG0539 | RpsA | Ribosomal protein S1 | Ribosome 30S subunit |
| COG0560 | SerB | Phosphoserine phosphatase | Serine biosynthesis |
| COG0578 | GlpA | Glycerol-3-phosphate dehydrogenase | Isoprenoid biosynthesis |
| COG0603 | QueC | 7-cyano-7-deazaguanine synthase (queuosine biosynthesis) | tRNA modification |
| COG0611 | ThiL | Thiamine monophosphate kinase | Thiamine biosynthesis |
| COG0643 | CheA | Chemotaxis protein histidine kinase CheA | |
| COG0659 | SUL1 | Sulfate permease or related transporter, MFS superfamily, contains STAS domain | |
| COG0685 | MetF | 5,10-methylenetetrahydrofolate reductase | |
| COG0686 | Ald | Alanine dehydrogenase (includes sporulation protein SpoVN) | Urea cycle |
| COG0687 | PotD | Spermidine/putrescine-binding periplasmic protein | |
| COG0688 | Psd | Phosphatidylserine decarboxylase | Phospholipid biosynthesis |
| COG0737 | UshA | 2',3'-cyclic-nucleotide 2'-phosphodiesterase/5'- or 3'-nucleotidase, 5'-nucleotidase family | |
| COG0738 | FucP | Fucose permease | |
| COG0757 | AroQ | 3-dehydroquinate dehydratase, type II | Aromatic amino acid biosynthesis |
| COG0759 | YidD | Membrane-anchored protein YidD, putatitve component of membrane protein insertase Oxa1/YidC/SpoIIIJ | |
| COG0770 | MurF | UDP-N-acetylmuramyl pentapeptide synthase | Mureine biosynthesis |
| COG0773 | MurC | UDP-N-acetylmuramate-alanine ligase MurC and related ligases, MurC/Mpl family | Mureine biosynthesis |
| COG0801 | FolK | 7,8-dihydro-6-hydroxymethylpterin pyrophosphokinase (folate biosynthesis) | Folate biosynthesis |
| COG0803 | ZnuA | ABC-type Zn uptake system ZnuABC, Zn-binding component ZnuA | |

| | | | |
|---|---|---|---|
| COG0826 | RlhA | 23S rRNA C2501 and tRNA U34 5'-hydroxylation protein RlhA/YrrN/YrrO, U32 peptidase family | 23S rRNA modification |
| COG0853 | PanD | Aspartate 1-decarboxylase | Pantothenate/CoA biosynthesis |
| COG0854 | PdxJ | Pyridoxine 5'-phosphate synthase PdxJ | Pyridoxal phosphate biosynthesis |
| COG0860 | AmiC | N-acetylmuramoyl-L-alanine amidase | |
| COG1008 | NuoM | NADH:ubiquinone oxidoreductase subunit 4 (chain M) | NADH dehydrogenase |
| COG1013 | PorB | Pyruvate:ferredoxin oxidoreductase or related 2-oxoacid:ferredoxin oxidoreductase, beta subunit | Pyruvate oxidation |
| COG1022 | FAA1 | Long-chain acyl-CoA synthetase (AMP-forming) | |
| COG1043 | LpxA | Acyl-[acyl carrier protein]--UDP-N-acetylglucosamine O-acyltransferase | Lipid A biosynthesis |
| COG1044 | LpxD | UDP-3-O-[3-hydroxymyristoyl] glucosamine N-acyltransferase | Lipid A biosynthesis |
| COG1047 | SlpA | Peptidyl-prolyl cis-trans isomerase, FKBP type | |
| COG1058 | CinA | ADP-ribose pyrophosphatase domain of DNA damage- and competence-inducible protein CinA | |
| COG1073 | FrsA | Fermentation-respiration switch esterase FrsA, DUF1100 family | |
| COG1089 | Gmd | GDP-D-mannose dehydratase | |
| COG1139 | LutB | L-lactate utilization protein LutB, contains a ferredoxin-type domain | |
| COG1151 | Hcp | Hydroxylamine reductase (hybrid-cluster protein) | |
| COG1166 | SpeA | Arginine decarboxylase (spermidine biosynthesis) | |
| COG1179 | TcdA | tRNA A37 threonylcarbamoyladenosine dehydratase | tRNA modification |
| COG1212 | KdsB | CMP-2-keto-3-deoxyoctulosonic acid synthetase | Lipid A biosynthesis |
| COG1230 | CzcD | Co/Zn/Cd efflux system component | |
| COG1270 | CbiB | Cobalamin biosynthesis protein CobD/CbiB | Cobalamine/B12 biosynthesis |
| COG1397 | DraG | ADP-ribosylglycohydrolase | |
| COG1435 | Tdk | Thymidine kinase | Pyrimidine salvage |
| COG1492 | CobQ | Cobyric acid synthase | Cobalamine/B12 biosynthesis |
| COG1530 | CafA | Ribonuclease G or E | |
| COG1558 | FlgC | Flagellar basal body rod protein FlgC | |
| COG1600 | QueG | Epoxyqueuosine reductase QueG (queuosine biosynthesis) | tRNA modification |
| COG1610 | YqeY | Uncharacterized conserved protein YqeY, may have tRNA amino acid amidase activity | |
| COG1619 | LdcA | Muramoyltetrapeptide carboxypeptidase LdcA (peptidoglycan recycling) | |
| COG1703 | ArgK | GTPase of the G3E family (not a periplasmic protein kinase) | |
| COG1726 | NqrA | Na+-transporting NADH:ubiquinone oxidoreductase, subunit NqrA | Na+-translocating NADH dehydrogenase |
| COG1778 | KdsC | 3-deoxy-D-manno-octulosonate 8-phosphate phosphatase KdsC and related HAD superfamily phosphatases | |
| COG1781 | PyrI | Aspartate carbamoyltransferase, regulatory subunit | |
| COG1785 | PhoA | Alkaline phosphatase | Folate biosynthesis |
| COG1805 | NqrB | Na+-transporting NADH:ubiquinone oxidoreductase, subunit NqrB | Na+-translocating NADH dehydrogenase |
| COG1838 | FumA | Tartrate dehydratase beta subunit/Fumarate hydratase class I, C-terminal domain | TCA cycle |
| COG1869 | RbsD | D-ribose pyranose/furanose isomerase RbsD | |
| COG2038 | CobT | NaMN:DMB phosphoribosyltransferase | Cobalamine/B12 biosynthesis |
| COG2066 | GlsA | Glutaminase | |
| COG2096 | PduO | Cob(II)alamin adenosyltransferase | Cobalamine/B12 biosynthesis |

| | | | |
|---|---|---|---|
| COG2152 | | Predicted glycosyl hydrolase, GH43/DUF377 family | |
| COG2160 | AraA | L-arabinose isomerase | |
| COG2200 | EAL | EAL domain, c-di-GMP-specific phosphodiesterase class I (or its enzymatically inactive variant) | |
| COG2209 | NqrE | Na+-transporting NADH:ubiquinone oxidoreductase, subunit NqrE | Na+-translocating NADH dehydrogenase |
| COG2226 | UbiE | Ubiquinone/menaquinone biosynthesis C-methylase UbiE/MenG or 23S rRNA G745 methylase RmlAII or tRNA-U5 methylase TRM9 | Biotin biosynthesis |
| COG2271 | UhpC | Sugar phosphate permease | |
| COG2310 | TerZ | Stress response protein SCP2 | |
| COG2382 | Fes | Enterochelin esterase or related enzyme | |
| COG2431 | LysO | Lysine export protein LysO/YbjE, DUF340 family | |
| COG2721 | UxaA | Altronate dehydratase | |
| COG2730 | BglC | Aryl-phospho-beta-D-glucosidase BglC, GH1 family | |
| COG2814 | AraJ | Predicted arabinose efflux permease AraJ, MFS family | |
| COG2829 | PldA | Outer membrane phospholipase A | |
| COG2871 | NqrF | Na+-transporting NADH:ubiquinone oxidoreductase, subunit NqrF | Na+-translocating NADH dehydrogenase |
| COG2877 | KdsA | 3-deoxy-D-manno-octulosonic acid (KDO) 8-phosphate synthase | Lipid A biosynthesis |
| COG2957 | AguA | Agmatine/peptidylarginine deiminase | |
| COG2963 | InsE | Transposase InsE and inactivated derivatives | |
| COG3004 | NhaA | Na+/H+ antiporter NhaA | |
| COG3033 | TnaA | Tryptophanase | |
| COG3041 | YafQ | mRNA-degrading endonuclease YafQ (mRNA interferase), toxin component of the YafQ-DinJ toxin-antitoxin module | |
| COG3075 | GlpB | Anaerobic glycerol-3-phosphate dehydrogenase | |
| COG3201 | PnuC | Nicotinamide riboside transporter PnuC | |
| COG3263 | NhaP2 | NhaP-type Na+/H+ and K+/H+ antiporter with C-terminal TrkAC and CorC domains | |
| COG3437 | RpfG | Response regulator c-di-GMP phosphodiesterase, RpfG family, contains REC and HD-GYP domains | |
| COG3661 | AguA2 | Alpha-glucuronidase | |
| COG3693 | XynA | Endo-1,4-beta-xylanase, GH35 family | |
| COG4108 | PrfC | Peptide chain release factor RF-3 | Translation factors |
| COG4124 | ManB2 | Beta-mannanase | |
| COG4591 | LolC | ABC-type lipoprotein targeting system transmembrane component LolC/LolE | |
| COG4659 | RnfG | Na+-translocating ferredoxin:NAD+ oxidoreductase RNF, RnfG subunit | Na+-translocating Fd:NADH oxidoreductase |
| COG4786 | FlgG | Flagellar basal body rod protein FlgG | |
| COG0419 | SbcC | DNA repair exonuclease SbcCD ATPase subunit | |
| COG0612 | PqqL | Predicted Zn-dependent peptidase, M16 family | |
| COG1228 | HutI | Imidazolonepropionase or related amidohydrolase | |
| COG1668 | NatB | ABC-type Na+ efflux pump, permease component NatB | |
| COG1690 | RtcB | RNA-splicing ligase RtcB, repairs tRNA damage | |
| COG2116 | FocA | Formate/nitrite transporter FocA, FNT family | |
| COG0131 | HisB2 | Imidazoleglycerol phosphate dehydratase HisB | Histidine biosynthesis |
| COG0459 | GroEL | Chaperonin GroEL (HSP60 family) | |

| | | | |
|---|---|---|---|
| COG0515 | SPS1 | Serine/threonine protein kinase, uncludes type III secretion system effector YopO | |
| COG0849 | FtsA | Cell division ATPase FtsA | |
| COG1167 | ARO8 | DNA-binding transcriptional regulator, MocR family, contains an aminotransferase domain | Lysine biosynthesis |

**Supplementary Table 7. Gene-body DMR genes associated with Factor 2.** Genes containing differentially methylated regions (DMRs) within their gene bodies that are associated with Factor 2 (n = 16).

| Ensemb ID | Entrez ID | Symbol | Gene Name | Chromosome | Microbiota Regulated | Transcription Factor |
|---|---|---|---|---|---|---|
| ENSMUSG00000034220 | 14733 | Gpc1 | glypican 1 | chr1 | Yes | No |
| ENSMUSG00000028655 | 76574 | Mfsd2a | MFSD2 lysolipid transporter A, lysophospholipid | chr4 | Yes | No |
| ENSMUSG00000036687 | 231832 | Tmem184a | transmembrane protein 184a | chr5 | Yes | No |
| ENSMUSG00000030214 | 66857 | Plbd1 | phospholipase B domain containing 1 | chr6 | Yes | No |
| ENSMUSG00000031561 | 23965 | Tenm3 | teneurin transmembrane protein 3 | chr8 | Yes | No |
| ENSMUSG00000010651 | 235674 | Acaa1b | acetyl-Coenzyme A acyltransferase 1B | chr9 | Yes | No |
| ENSMUSG00000000594 | 14667 | Gm2a | GM2 ganglioside activator protein | chr11 | Yes | No |
| ENSMUSG00000073460 | 240023 | Pnldc1 | poly(A)-specific ribonuclease (PARN)-like domain containing 1 | chr17 | Yes | No |
| ENSMUSG00000005373 | 58805 | Mlxipl | MLX interacting protein-like | chr5 | No | Yes |
| ENSMUSG00000087382 | 74161 | Ctcflos | CCCTC-binding factor like, opposite strand | chr2 | No | No |
| ENSMUSG00000028028 | 71481 | Alpk1 | alpha-kinase 1 | chr3 | No | No |
| ENSMUSG00000053898 | 51798 | Ech1 | enoyl coenzyme A hydratase 1, peroxisomal | chr7 | No | No |
| ENSMUSG00000020865 | 76408 | Abcc3 | ATP-binding cassette, sub-family C member 3 | chr11 | No | No |
| ENSMUSG00000025792 | 27376 | Slc25a10 | solute carrier family 25 (mitochondrial carrier, dicarboxylate transporter), member 10 | chr11 | No | No |
| ENSMUSG00000114710 | 105245474 | Gm40923 | predicted gene, 40923 | chr13 | No | No |
| ENSMUSG00000052133 | 20357 | Sema5b | sema domain, seven thrombospondin repeats (type 1 and type 1-like), transmembrane domain (TM) and short cytoplasmic domain, (semaphorin) 5B | chr16 | No | No |

**Supplementary Table 8.** Table of the monitored ion transitions from the epigenetically relevant metabolites.

| Metabolite | Formula | Monoisotopic Mass | Parent m/z | m/z 1st transition (CE) | m/z 2n transition (CE) | Polarity |
|---|---|---|---|---|---|---|
| Adenosine | $C_{10}H_{13}N_5O_4$ | 267.096 | 268 | 136 (16) | 119 (48) | Positive |
| Adenosine diphosphate (ADP) | $C_{10}H_{15}N_5O_{10}P_2$ | 427.029 | 428 | 136 (32) | 348 (16) | Positive |
| ADP-Ribose | $C_{15}H_{23}N_5O_{14}P_2$ | 559.071 | 560 | 136 (36) | 348 (16) | Positive |
| Ac-ADP-Ribose | $C_{17}H_{25}N_5O_{15}P_2$ | 601.082 | 602 | 136 (36) | 348 (16) | Positive |
| Adenosine Triphosphate (ATP) | $C_{10}H_{16}N_5O_{13}P_3$ | 506.996 | 508 | 136 (44) | 97 (40) | Positive |
| Uridine diphosphate (UDP) | $C_9H_{14}N_2O_{12}P_2$ | 404.002 | 405 | 97 (24) | 113 (36) | Positive |
| UDP-Glucose-N-Acetylglucosamine (UDP-Glc-Nac) | $C_{17}H_{27}N_3O_{17}P_2$ | 607.082 | 608 | 204 (8) | 138(48) | Positive |
| S-Adenosyl Methionine (SAM) | $C_{15}H_{22}N_6O_5S$ | 399.145 | 399 | 250 (12) | 97 (32) | Positive |
| S-Adenosyl-Homocysteine (SAH) | $C_{14}H_{20}N_6O_5S$ | 384.122 | 385 | 136 (24) | 250 (4) | Positive |
| Betaine | $C_5H_{11}NO_2$ | 117.079 | 118 | 58(28) | 59(16) | Positive |
| Glutamine | $C_5H_{10}N_2O_3$ | 146.069 | 147 | 84 (20) | 130 (8) | Positive |

| | | | | | | |
|---|---|---|---|---|---|---|
| Histidine | $C_6H_9N_3O_2$ | 155.069 | 156 | 110 (12) | 83 (28) | Positive |
| Methionine | $C_5H_{11}NO_2S$ | 149.051 | 150 | 56 (12) | 104 (6) | Positive |
| Serine | $C_3H_7NO_3$ | 105.042 | 106 | 60 (12) | 42 (20) | Positive |
| Dimethylglycine | $C_4H_9NO_2$ | 103.063 | 104 | 58 (12) | 42 (35) | Positive |
| α-Ketoglutarate | $C_5H_6O_5$ | 144.006 | -145 | -101 (4) | -57 (8) | Negative |
| Citrate | $C_6H_8O_7$ | 189.004 | -191 | -67 (28) | -57(20) | Negative |
| Succinate | $C_4H_6O_4$ | 118.026 | -117 | -73 (8) | -99 (8) | Negative |
| Glucose | $C_6H_{12}O_6$ | 180.063 | 181 | 99 (12) | 140 (4) | Positive |
| Choline | $C_5H_{14}NO$ | 104.108 | 104 | 60 (20) | 45 (15) | Positive |
| Acetyl-CoA | $C_{23}H_{38}N_7O_{17}P_3S$ | 809.126 | 810 | 303 (36) | 136 (60) | Positive |
| Coenzyme-A | $C_{21}H_{36}N_7O_{16}P_3S$ | 767.115 | 768 | 261 (30) | 428 (25) | Positive |
| 5-Methyl-Tetrahydrofolate | $C_{20}H_{25}N_7O_6$ | 459.187 | 460 | 313 (24) | 180 (44) | Positive |
| Oxidized Nicotinamide adenine dinucleotide ($NAD^+$) | $C_{21}H_{28}N_7O_{14}P_2^+$ | 664.117 | 664 | 136 (60) | 428 (24) | Positive |
| Reduced NAD (NADH) | $C_{21}H_{29}N_7O_{14}P_2$ | 665.125 | 666 | 136 (40) | 137 (52) | Positive |
| Flavin adenine dinucleotide (FAD) | $C_{27}H_{33}N_9O_{15}P_2$ | 785.157 | 786 | 348 (24) | 136 (48) | Positive |
| Riboflavin | $C_{17}H_{20}N_4O_6$ | 376.138 | 377 | 243 (20) | 172 (35) | Positive |
